# Distinct functions of *PBRM1* and *BAP1* reconcile the course of kidney cancer evolution and disease progression

**DOI:** 10.64898/2026.04.30.721837

**Authors:** Daqi Deng, Hugang Feng, Annika Fendler, Yuliia Dovga, Fiona Byrne, Charlotte Spencer, Alice C. Martin, Ángel Fernández Sanromán, Omar Bouricha, Scott T. C. Shepherd, Hongchang Fu, Anne-Laure Cattin, Husayn Pallikonda, Irene Lobon, Kevin Mulder, Qingli Guo, Matouš Elphick, Jiahao Wang, Antonia Franz, Suzan Ben-Akinduro, Zayd Tippu, Barbara Ibarzo Yus, Taja Barber, Stephanie Hepworth, Isabel Symons, Kim Edmonds, Eleanor Carlyle, Arjun Modi, Justine Korteweg, Andressa Schneiders Santos, Leo Bickley, Sarah Rudman, Axel Bex, James Larkin, Samra Turajlic

## Abstract

Clear cell renal cell carcinoma (ccRCC) progresses along two predominant evolutionary trajectories, defined by *PBRM1* (∼40%) or *BAP1* (∼15%) mutations on a *VHL*-inactivated background. They have distinct patterns of evolutionary tempo and mode, and vastly different clinical outcomes, yet the underlying genotype-specific molecular phenotypic programmes are unknown. We established a patient-derived preclinical model biobank that captures the genetic diversity of ccRCC. Through integrative analyses of preclinical models and tumour bulk and single cell profiling, we identified transcriptional and epigenetic changes specific to *PBRM1*- and *BAP1*-driven ccRCC. Modelling *PBRM1* loss *in vitro* demonstrates that it reinforces renal lineage identity and maintains progenitor-like cell state. In contrast, *BAP1* loss drives inflammatory signalling and chromosomal instability. These insights reconcile the distinct evolutionary modes (branched versus punctuated), tempo (slow versus fast) and clinical outcomes associated with *PBRM1* and *BAP1* mutations, respectively, establishing a framework for patient stratification and genotype-directed therapeutic development.

## INTRODUCTION

Kidney cancer is a significant health burden globally, and there is an increase in both incidence and mortality rates^1,2^. ccRCC is the predominant histological subtype, representing approximately 75% of all cases^3^. ccRCC is characterised by stereotyped genetics. Early ccRCC development proceeds through the loss of the short arm of chromosome 3 (3p) and inactivation of the Von Hippel-Lindau (*VHL*) tumour suppressor gene, observed in >90% of all cases^4–8^. Tumours harbouring these events alone are often indolent and associated with an early disease stage^8,9^. However, the majority of ccRCC evolve beyond these initial events, acquiring additional genetic drivers and phenotypes, and it is the disease’s genetic progression that drives a more aggressive clinical phenotype. This clinical reality underscores a critical need to deepen our mechanistic understanding of ccRCC progression to inform rational therapeutic strategies and improve disease management.

ccRCC progresses through the acquisition of additional genetic alterations and dynamic interactions with the tumour microenvironment^10^. Large-scale genomic studies have identified frequent mutations in chromatin-remodelling genes located on chromosome 3p, particularly Polybromo 1 (*PBRM1*) and BRCA1-associated protein 1 (*BAP1*)^11–14^. These alterations act as secondary drivers that define distinct evolutionary trajectories and are associated with divergent clinical behaviours^15,16^. *PBRM1*, a component of the PBAF chromatin remodelling complex, plays a central role in regulating chromatin accessibility and transcription^17,18^. Primary tumours with *PBRM1* mutations develop over long periods of time, are characterised by increasing subclonal diversification and a branched evolutionary pattern typically associated with a more indolent disease behaviour^9,19^. *BAP1* encodes a deubiquitinase and regulates transcription through histone modification^20–22^. *BAP1*-mutated tumours evolve rapidly following a punctuated evolutionary trajectory leading to dominance of a highly fit clone and low intratumoural heterogeneity, often exhibiting high aneuploidy level, increased proliferative capacity, and a markedly aggressive clinical phenotype^16,23,24^.

Although these two types of ccRCC have been recognised for over a decade, the distinct mechanisms by which they drive tumour progression remain elusive, and genotype-specific therapeutic strategies are lacking. In this study, we leveraged clinical specimens from the TRACERx Renal study to establish a biobank of patient-derived preclinical models representing diverse genomic backgrounds. Transcriptomic and epigenomic profiling revealed distinct molecular features in *BAP1*- and *PBRM1*-mutant preclinical models. Integrative analysis of publicly available bulk and single-cell RNA-seq datasets, together with loss-of-function modelling using CRISPR/Cas9-engineered organoids, further supported the existence of two dominant onco-programmes. *PBRM1* loss promotes a permissive state, characterised by renal lineage retention, enhanced hypoxia signalling, and fibrotic tumour microenvironment (TME), which collectively facilitate sustained progression. In contrast, loss of *BAP1* drives an aggressive phenotype by inducing chromosomal instability (CIN), and *BAP1*-mutant tumours were characterised by high proliferative and stress response signatures, producing an inflamed TME.

## RESULTS

### A ccRCC preclinical model biobank

Functional understanding of ccRCC is hindered by the shortage of representative *in vitro* models^25,26^, particularly those capturing its genomic diversity. With preclinical model derivation attempts from 55 patients recruited through the TRACERx Renal project^9^ (102 tumour and 42 normal kidney specimens), we established 18 preclinical cancer models, comprising 11 organoids, two spheroids, and five 2D cell lines, as well as 42 normal kidney organoids (**Figure 1A**, **Figure S1A**, **Table S1**). Normal kidney organoids typically formed cystic or tubular structures, while tumour-derived organoids exhibited solid and cohesive or rosette-like and non-cohesive morphologies (**Figure 1B**). Some cultures also exhibited a spread-out, mesenchymal-like growth pattern. Histologically, tumour organoids retained key features of primary tumours, including clear cytoplasm and irregular nuclear morphology (**Figure 1B**). Notably, all models derived from early-stage (Stage I and II) tumours failed to expand beyond the initial passages, despite maintaining viable morphology and cell-cell contacts. These cultures exhibited cell cycle arrest rather than apoptosis (data not shown), suggesting an intrinsic barrier to *in vitro* propagation. In contrast, organoids established from advanced-stage tumours demonstrated sustained expansion in culture (**Table S1**). This observation aligns with findings from traditional 2D cell line models, which predominantly reflect a more aggressive disease state^26^.

**Figure 1.**
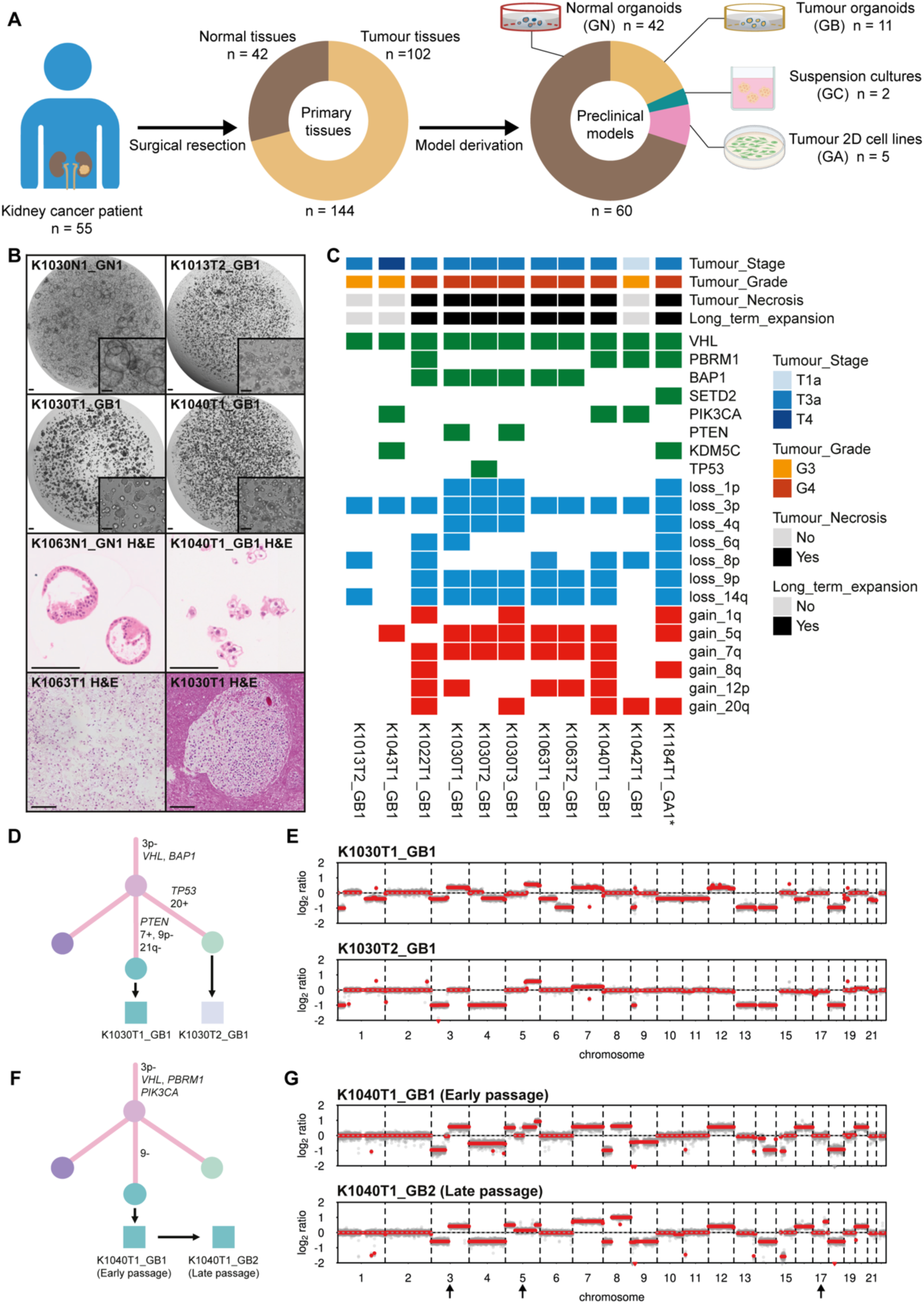
A patient-derived preclinical model biobank recapitulates the genomic diversity of ccRCC. **(A)** Overview of ccRCC preclinical model derivation. (B) Representative bright field images (top), H&E staining of organoids (mid) and primary tumours (bottom). Scale bar, 100 μm. (C) Genomic landscape of patient-derived preclinical models, highlighting key driver mutations across the cohort. *: treated sample. (D) Phylogenetic reconstruction of case K1030 showing clonal relationships between primary tumour clones (circles) and derived models (squares). (E) Comparison of genome-wide copy number profiles from two independent organoid lines (K1030T1_GB1 and K1030T2_GB1) derived from distinct tumour regions of patient K1030. (F) Phylogenetic reconstruction of case K1040 showing clonal relationships between primary tumour clones (circles) and corresponding organoid models (squares). (G) Comparison of copy number profiles from a longitudinally cultured organoid line (from K1040T1_GB1 to K1040T1_GB2), illustrating ongoing karyotypic evolution *in vitro*. See also Figure S1-2 and Table S1.

To evaluate how well the organoids recapitulate the genomic landscape of ccRCC, we performed genomic profiling using a targeted ccRCC driver panel sequencing approach as described before^9^. We selected 10 patient-derived organoids (PDOs) and one patient-derived cell line (PDCL) for genomic profiling. We also examined *VHL* promoter methylation to identify *VHL* inactivation via epigenetic mechanisms. All models harboured 3p loss and *VHL* inactivation, consistent with their prevalence in clinical sample cohorts^8,12^. *VHL* inactivation in K1022T1_GB1 occurred via promoter methylation, while K1030-derived organoids harboured a novel intronic mutation disrupting splicing (**Figure S1B**), highlighting diverse inactivation mechanisms underlying the selection of *VHL* loss. Long-term propagating models exhibited extensive copy number variations (CNVs), including losses of 9p and 14q (**Figure 1C**). These profiled preclinical models captured the major ccRCC genotypes. K1013T2_GB1 harboured minimal alterations, including *VHL* mutation, 3p loss, 8p loss, and 14q loss. K1030T1_GB1, K1030T2_GB1, K1030T3_GB1, K1063T1_GB1, and K1063T2_GB1 were *BAP1* driven, while K1040T1_GB1 and K1042T1_GB1, K1184T1_GA1 were *PBRM1*-deficient with additional *SETD2* or *PIK3CA* mutations. K1022T1_GB1 captured a scenario in which *BAP1* and *PBRM1* co-occurred, representing a rare multiple-clonal driver subtype (**Figure 1C**).

Organoid cultures captured intratumor heterogeneity (ITH) and showed *ex vivo* evolution. Multiregional genomic profiling of the K1030 primary tumour revealed subclonal *PTEN* and *TP53* mutations in the background of clonal 3p loss, *VHL* mutation, and *BAP1* mutation. As a result of multiregional PDO derivation, K1030T1_GB1 captured the “*VHL* → *BAP1* → *PTEN*” subclone, while K1030T2_GB1 captured the “*VHL* → *BAP1* → *TP53*” subclone. These lines also differed in their CNV profiles, with K1030T1_GB1 exhibiting a higher burden of somatic copy number alterations (SCNAs) (**Figure 1D-E**). Although the long-term cultured organoids were derived from late-stage tumours, ongoing *in vitro* evolution was also observed. For example, in K1040T1_GB1, additional chromosomal alterations, including more variations in Chr 3, 5, and 17, emerged over passages (**Figure 1F-G**). We considered the evolved organoid as a new line (K1040T1_GB2). Overall, these characterisations highlight that this preclinical model cohort represents the genomic diversity and evolutionary features of ccRCC.

### BAP1 mutants exhibit higher proliferative capacity

To characterise the transcriptomic landscape of the preclinical models, we profiled single-cell transcriptomes from a subset of cultures. After quality control and filtering, 25,180 cells were retained, representing 11 tumour lines and one normal line (**Figure 2A**, **Table S2**). The ccRCC marker caeruloplasmin (*CP*) was expressed in all tumour organoid models but not in the tumour cell line (K1184T1_GA1) and normal organoid (K1030N1_GN1) (**Figure 2B**). Consistent with genomic sequencing data, inferCNV revealed that long-term expanded organoids exhibited extensive CNVs (**Figure S2**).

**Figure 2.**
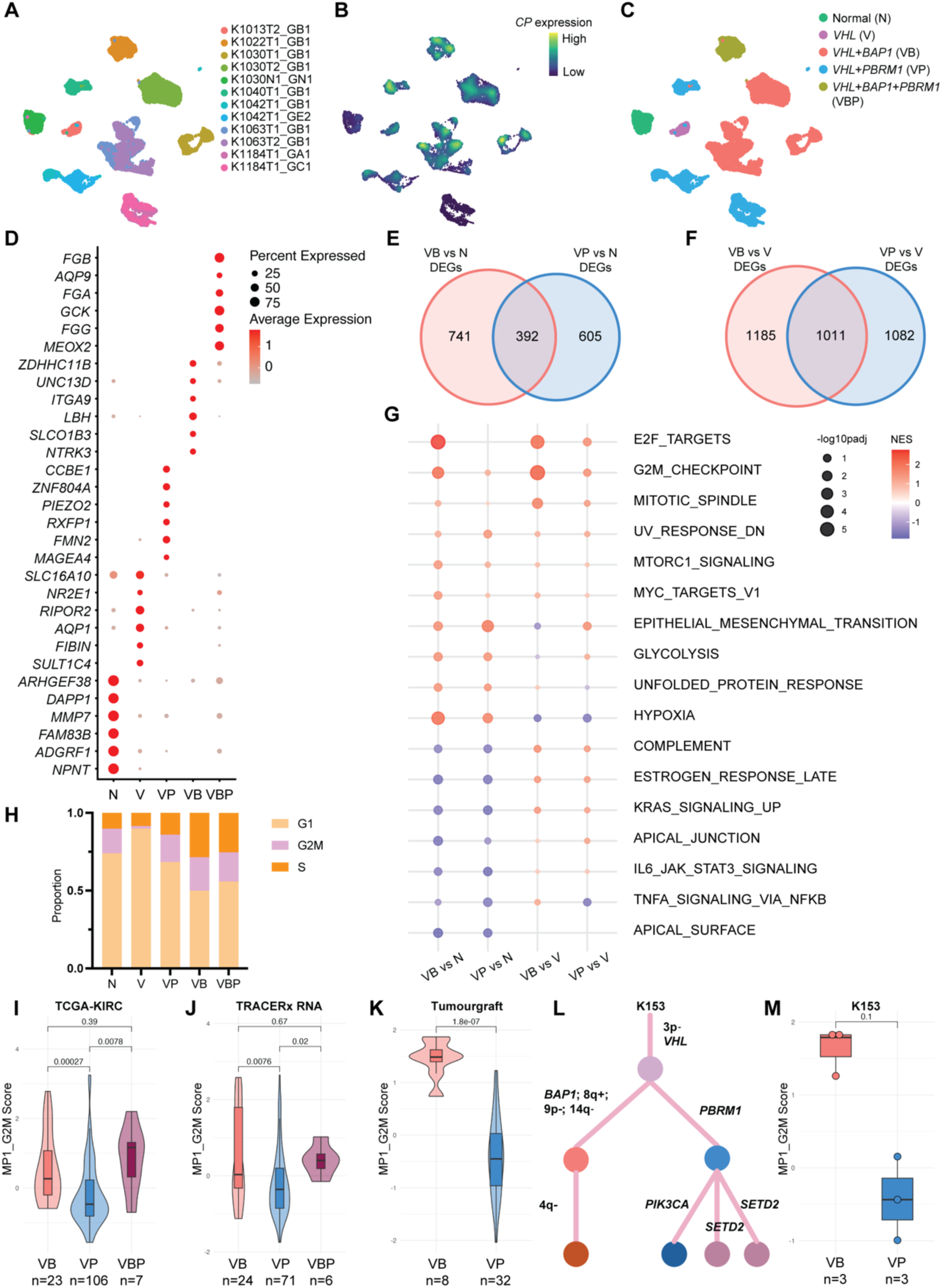
Transcriptional characterisations of ccRCC organoids with *BAP1* and *PBRM1* mutations. **(A-C)** Uniform manifold approximation and projection (UMAP) of single cell transcriptomes from ccRCC preclinical models, coloured by (A) line identity, (B) expression level of *Ceruloplasmin* (*CP*), and (C) mutation status. N: normal, V: *VHL*, VB: *VHL*+*BAP1*, VP: *VHL*+*PBRM1*, VBP: *VHL*+*BAP1*+*PBRM1*. (D) Dot plot showing the top 6 differentially upregulated genes in five genotypes. Dot size represents the proportion of cells expressing each gene, and colour intensity reflects the mean normalised expression level. (E and F) Numbers of unique and shared upregulated genes in *BAP1*- and *PBRM1*-mutant organoids (VB and VP, respectively) compared to (E) normal organoids and (F) *VHL*-only organoids. (G) Normalised enrichment scores (NES) for Hallmark pathways based on differentially expressed genes between VB or VP organoids and normal or *VHL*-only organoids. Dot size denotes pathway significance, and colour gradient represents NES (red, enriched; blue, depleted). (H) Comparison of cell cycle phase distribution across organoids with different genotypes. (I-K) Violin plots showing proliferation capability (G2M meta program as a surrogate) difference across *BAP1*-driven and *PBRM1*-driven tumours in (I) TCGA-KIRC cohort, (J) TRACERx RNA cohort, and (K) Tumourgraft cohort. Sample sizes are denoted under the group names. (L) Phylogenetic tree of case K153 showing two independent branches (*BAP1*-driven and *PBRM1*-driven) originating from the same *VHL* loss trunk. (M) Box plot showing proliferation capability difference between *BAP1*-driven and *PBRM1*-driven branches in case K153. Sample sizes are denoted under the group names. See also Figure S3-4, Table S2-4.

*PBRM1* and *BAP1* are two major secondary drivers in 40% and 15% of ccRCC cases, respectively^9^. Therefore, we grouped the preclinical models by genotype to investigate genotype-phenotype relationships (**Figure 2C**). Differentially expressed analysis revealed distinct transcriptional patterns across genotypes (**Figure 2D**). We compared *PBRM1*- or *BAP1*-driven subtypes (referred to as VP or VB, respectively) to the normal renal organoids (referred to as N). There were 392 upregulated genes shared between both subtypes, whereas 741 genes were *BAP1*-specific and 605 genes were *PBRM1*-specific (log2 fold change (LFC) > 0.5 & padj < 0.05, **Figure 2E, Table S3**). Comparing these two subtypes to their ancestor genotype *VHL* only (referred to as V) revealed 1185 genes uniquely upregulated in *BAP1*-driven organoids, 1082 in *PBRM1*-driven organoids, and 1011 genes shared between both groups (LFC > 0.5 & padj < 0.05, **Figure 2F**, **Table S3**). Gene set enrichment analysis (GSEA) showed hypoxia and glycolysis were significantly upregulated across tumour organoids, along with epithelial-mesenchymal transition (EMT) and unfolded protein response (**Figure 2G**). While *PBRM1*-driven organoids did not exhibit increased proliferative capacity compared to normal organoids, as indicated by the absence of enrichment in proliferation-related pathways, *BAP1*-driven tumour organoids displayed strong upregulation of cell cycle hallmarks, including E2F targets and G2M checkpoint genes, reflecting their more aggressive phenotype (**Figure 2G**). Inference of cell cycle phase from scRNA-seq data also showed a higher proportion of G2M and S phase cells in the *BAP1* group (**Figure 2H**).

To validate the proliferation capacity difference observed between *BAP1* and *PBRM1* mutants, we collected and analysed bulk RNA-seq data from three independent ccRCC cohorts: TCGA-KIRC^12^, TRACERx Renal^27^, and a ccRCC tumour xenograft (referred to as tumourgraft, TG) cohort^28^ (**Table S4**). We derived a transcriptional signature landscape of clinical specimens by calculating the single-sample GSEA (ssGSEA) scores of 39 gene sets, including hallmark gene sets, scRNA-seq-derived meta-programmes (MPs), and publication-based gene signatures (**Table S4**, **Figure S3-S4**, **Methods**). As expected, G2M metaprogram, as an indicator of proliferation capacity, was significantly higher in *BAP1* mutants compared to *PBRM1* mutants in all three cohorts (**Figure 2I-K**, **Figure S3-S4**).

A case study of a rare scenario where *PBRM1* and *BAP1* independently arise in the same tumour, forming two distinct subclones, offers valuable insights into the functional divergence between these drivers. Multiregional panel sequencing of a unique case (K153) from the TRACERx Renal study revealed a tumour harbouring two evolutionary branches^9^, both originating from a common ancestor with 3p loss and *VHL* mutation. One branch was driven by *BAP1* loss, while the other was defined by *PBRM1* loss (**Figure 2L**). The *BAP1*-mutant subclone showed a trend toward higher proliferation capacity, though this was not significant due to the small sample size (**Figure 2M**). Overall, these data from organoid and clinical specimen profiling demonstrated that *BAP1*-driven ccRCC are more proliferative.

### Intrinsic hypoxia response signature in PBRM1-mutated tumour cells

*PBRM1* mutation has been linked to hypoxia response amplification^29^. However, when we examined the hypoxia-related signatures across the three bulk RNA-seq cohorts, this difference was not significant (**Figure S5A**). The hypoxia response difference was also not observed in preclinical models (**Figure 2G**), presumably because all tumour models were cultured under the same hypoxic condition. The bulk measurement of hypoxia response would be affected by tumour microenvironment cells under hypoxic conditions^30^, which could lead to this discrepancy. Therefore, we thought to examine tumour intrinsic features at single-cell resolution by integrating six publicly available scRNA-seq datasets (a total of 44 tumour samples with genotype data available)^31–36^, as well as our preclinical model scRNA-seq dataset. For clinical tumour samples, we applied inferCNV to call chromosome 3p loss, and cells with 3p loss were identified as high-confidence tumour cells for downstream analysis (**Figure 3A, Methods**). This resulted in a total of 119,036 high-confidence tumour cells (mean = 2,705; range = 2-27,369) (**Figure S5B**). We grouped cells based on the presence of mutations, including “*VHL*”, “*BAP1*_any”, “*PBRM1*_any”, and “*BAP1*_*PBRM1*”. Compared to *VHL*-only tumour cells, *PBRM1*_any cells showed significant enrichment of the hypoxia pathway, with upregulation of canonical HIF-regulated genes like *TMEM45A* and *ANGPTL4* (**Figure 3B**). Direct comparison of *BAP1*_any and *PBRM1*_any groups showed that *PBRM1*-mutant cells exhibited upregulation of several hypoxia-related genes, including *HIF1A*, *EPAS1*, *VEGFA*, *TMEM45A*, and *EPO* (**Figure 3C**). Examination of the HIF metagene score also showed higher HIF response in *PBRM1* mutants (**Figure 3D**), further supporting the notion that hypoxia signalling is an intrinsic hallmark of *PBRM1*-mutant tumour cells compared to *BAP1*-mutant cells at the single-cell level.

**Figure 3.**
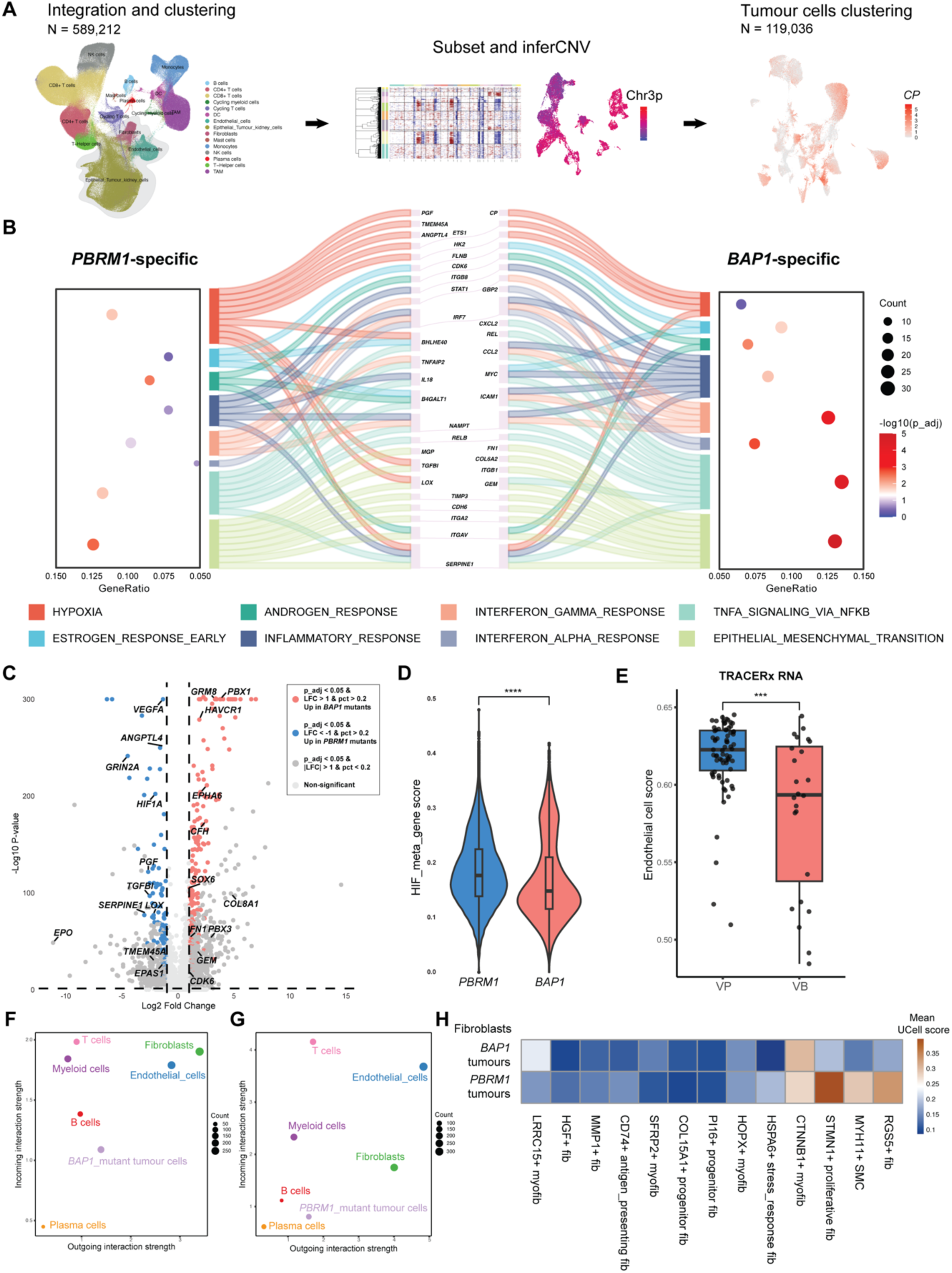
Intrinsic and extrinsic programmes associated with *BAP1* and *PBRM1* mutations. **(A)** Schematic overview of the curation and integration workflow for publicly available single-cell RNA-seq datasets. Single cells from six studies were integrated and annotated, followed by inferCNV-based identification of tumour cells, which were subsequently subset for differential expression analyses (see also *Methods*). (B) Mirrored Sankey plot showing hallmark pathway enrichment derived from differentially upregulated genes in *BAP1*- and *PBRM1*-mutant tumours relative to *VHL*-monodriver tumours. Pathways enriched in *PBRM1*-mutant tumours are shown on the left, and those enriched in *BAP1*-mutant tumours are on the right. The central section displays individual genes contributing to these pathways, with shared genes underlined and connected to multiple pathways. (C) Volcano plot comparing single-cell transcriptomes of *BAP1*-mutant and *PBRM1*-mutant tumour cells. Differentially expressed genes (adjusted *p* < 0.01, LFC > 1, expressed in >20% of cells) are highlighted in dark red (*BAP1*-specific) or dark blue (*PBRM1*-specific). (D) Violin plot comparing HIF meta-gene *UCell* scores between *BAP1*-mutant and *PBRM1*-mutant tumour cells (Wilcoxon signed-rank test, **** *p* < 0.0001). (E) Box plot comparing endothelial cell score between *BAP1*-driven and *PBRM1*-driven tumours in TRACERx RNA cohort (VB: n=24, VP: n=71, Wilcoxon signed-rank test, *** *p* < 0.001). (F-G) Scatter plots showing the signalling roles of different cell types in (F) *BAP1*-driven tumours and (G) *PBRM1*-driven tumours. The x-axis and y-axis represent the probabilities of outgoing and incoming communication, respectively. Dot size indicates the number of inferred links (both outgoing and incoming). Cell types are labelled and coloured. (H) Heatmap displaying mean *UCell* scores as a proxy for similarity between known fibroblast subtypes and fibroblasts derived from *PBRM1*- and *BAP1*-mutant tumours. See also Figure S5 and Table S4-5.

### Differential tumour microenvironmental remodelling in PBRM1- and BAP1-driven tumours

Tumour vasculature is associated with the level of hypoxia response of tumour cells^37^. Therefore, we further examined endothelial composition in *PBRM1*- and *BAP1*-driven tumours using ConsensusTME deconvolution^38^ (**Figure S5C, Methods**). As expected, *PBRM1*-driven tumours have a significantly higher endothelial score in the TRACERx RNA cohort (**Figure 3E**). However, in the TCGA-KIRC cohort, there is a non-significant trend (**Figure S5D**). Together, single-cell and bulk RNA-seq data suggest that *PBRM1*-mutated tumour cells intrinsically upregulate the hypoxia response, which may contribute to endothelial recruitment and vascularisation.

We further explored the tumour microenvironmental remodelling of *BAP1*- and *PBRM1*-driven tumours. *BAP1*-specific tumour cells showed strong enrichment for general inflammatory responses and chemokine signalling, with high expression *of CCL2*, *ICAM1*, and *CXCL2* (**Figure 3B**), acting as an active driver in the TME. Cell-cell communication analysis also showed that, in *BAP1*-driven tumours, incoming interactions in immune cells, especially myeloid cells, were stronger than in *PBRM1*-driven tumours (**Figure 3F-G**). In contrast, *PBRM1* mutant cells are more passive in the tumour microenvironment (**Figure 3G**).

The different strengths of fibroblast-centred communication networks in the two tumour types also prompted us to examine their fibroblast phenotypes (**Figure 3F-G**). We adapted the signatures of 13 non-tissue-specific fibroblast subtypes^39^ and analysed the similarity to fibroblasts in *PBRM1*-mutated tumours or *BAP1*-mutated tumours. Fibroblasts in *BAP1* tumours were enriched with LRRC15+ myofibroblasts, which were linked to high TGF-β signalling activity and tumour epithelial-to-mesenchymal transition (EMT) process^39,40^. Fibroblasts in *PBRM1* tumours were enriched with RGS5+ fibroblasts, MYH11+ smooth muscle cells, and STMN1+ proliferative fibroblasts (**Figure 3H**). These specialised fibroblast subtypes are associated with proliferation, stress response, and vasculature^39^. This suggests micro niches supporting tumour cell expansion in *PBRM1*-driven tumours. Additionally, in *PBRM1*-mutant cells, Matrix Gla Protein (*MGP*), TGF-β-induced protein (*TGFBI*), and Lysyl Oxidase (*LOX*) are differentially upregulated (**Figure 3B**). These genes are associated with extracellular matrix (ECM) remodelling, particularly fibrosis formation^41–43^. Indeed, during primary tissue processing, we observed that bulk tumours with *PBRM1* mutations showed a higher prevalence of fibrosis. VHL patients harbour a germline *VHL* mutation and are at risk of developing multiple clonally independent tumours in the kidney. Histological examination of the two independent tumours (one harbouring a *BAP1* mutation and the other harbouring a *PBRM1* mutation) arising in the same VHL patient also supports the notion of a *PBRM1*-specific microenvironment characterised by sclerotic, nested, and large stromal areas (**Figure S5E-F**). Together, these data suggest *BAP1*- and *PBRM1*-tumours exhibit distinct TME remodelling. While *BAP1*-driven tumours are more inflamed and have more interactions with TME cells, *PBRM1*-driven tumours are particularly enriched with endothelial cells and tend to form fibrosis in the tumour bed.

### Epigenetic landscapes in PBRM1- and BAP1-driven tumour cells

Both *PBRM1* and *BAP1* are chromatin remodellers^44^, and their inactivation could lead to reprogramming of the epigenetic landscape. Therefore, we applied SCENIC^45^, a tool commonly used in scRNA-seq data to infer regulon (a regulatory hub defined by a transcription factor (TF) and its target genes) activity, to explore the epigenetic regulation landscape in preclinical models (**Methods**). This revealed 299 regulons to be active in the preclinical model scRNA-seq data (**Table S2**). UMAP clustering of the regulon activity revealed that the normal organoid (K1030N1_GN1) formed a distinct cluster characterised by high activities in ELF1, ELF5, GATA3, and REL (**Figure 4A-B**), which are known marker TFs in proximal tubule cells^46–48^. Interestingly, the *VHL* organoid K1013T2_GB1 (V) clustered with K1042T1_GB1 (VP), which was *PBRM1*-driven, but at an early stage (**Figure 4A**). Less characterised TFs, such as VSX1 and RUNX3, were highly active in *PBRM1*-driven models (**Figure 4B**, **Figure S6A**). In *BAP1*-driven models, the stress response TFs such as FOS and JUN were highly active (**Figure 4B**). This is consistent with a recent report showing increased accessibility of stress response TF binding sites in ccRCC with *BAP1* mutation via single-cell Assay for Transposase-Accessible Chromatin using sequencing (scATAC-seq)^49^. Moreover, STAT6, a known inflammatory TF^50^, were more active in *BAP1*-driven PDOs (**Figure 4B**), suggesting an inflammatory phenotype independent of the tumour microenvironment.

**Figure 4.**
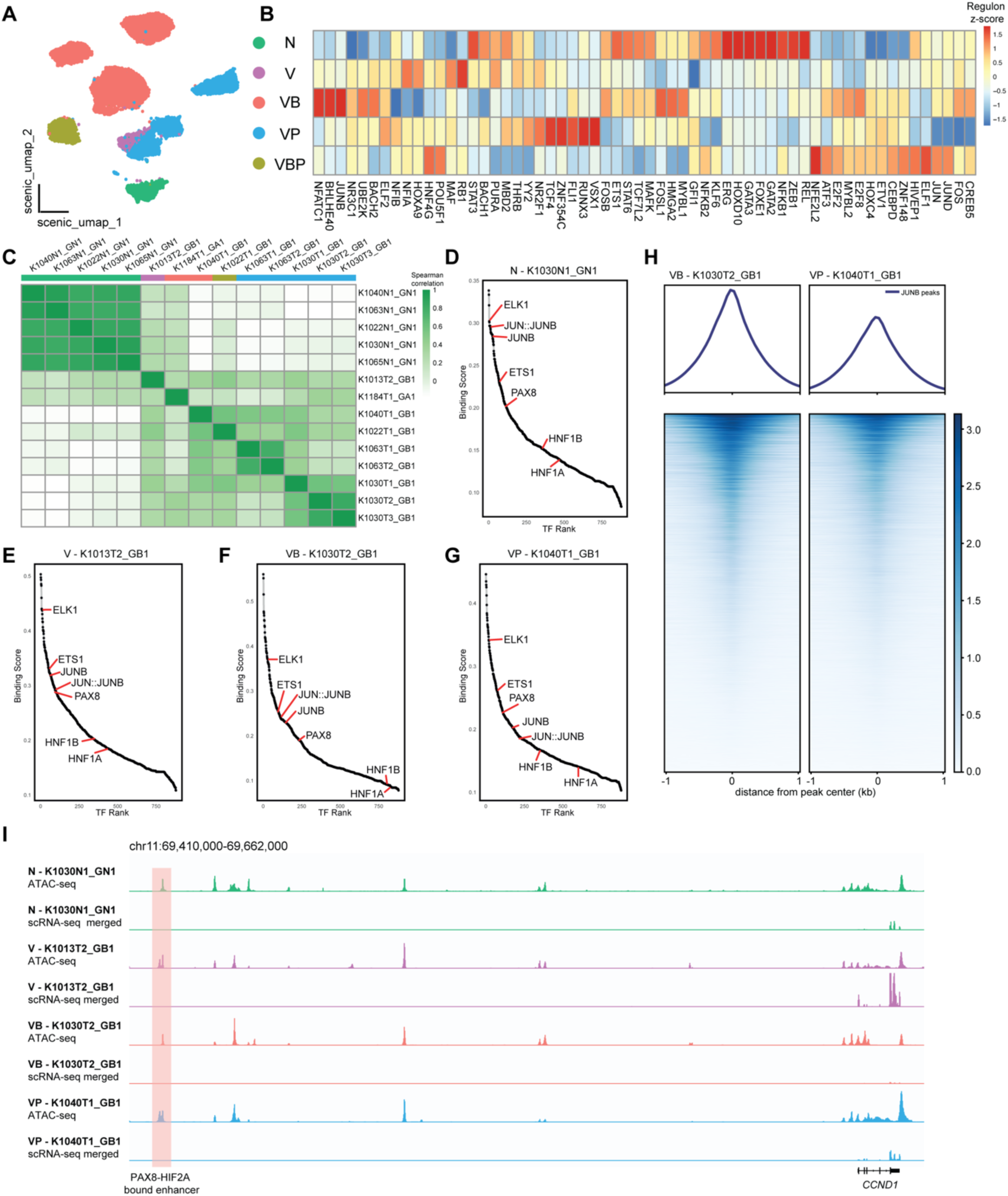
Epigenetic landscapes of ccRCC organoids with *BAP1* and *PBRM1* mutations. (A) UMAP of single-cell regulon activity from ccRCC organoids coloured by genotype. (B) Heatmap showing z-scores of top 60 variable regulons across organoids with different genotypes (genotype annotation as in panel A). (C) Heatmap displaying pairwise correlations between ATAC-seq samples based on the top 5,000 variable peaks. (D–G) Ranked transcription-factor (TF) footprinting plots for (D) normal organoid K1030N1_GN1, (E) *VHL* organoid K1013T2_GB1, (F) *BAP1*-driven organoid K1030T2_GB1, and (G) *PBRM1*-driven organoid K1040T1_GB1. Kidney-specific TFs (ELK1, ETS1, PAX8, HNF1A, HNF1B) and stress-response TFs (JUNB, JUN::JUNB) are annotated. (H) Heatmap and density distribution of ATAC-seq accessibility within ±1 kb of JUNB-binding peaks. (I) Coverage plots showing ATAC-seq accessibility and RNA-seq expression at the *CCND1* locus across normal (N), *VHL*-mutant (V), *BAP1*-mutant (VB), and *PBRM1*-mutant (VP) organoids. The shaded region marks the distal enhancer co-bound by PAX8 and HIF2A. See also Figure S6 and Table S2-3.

To further investigate the epigenetic regulation of *PBRM1* and *BAP1*-specific genes, we performed bulk ATAC-seq on organoids with distinct genotypes. Normal renal organoids showed a similar chromatin accessibility landscape, whereas tumour organoids showed patient-specific patterns (**Figure 4C**). We then performed TF footprinting analysis to identify specific TF activities across organoids. ETS-family TFs, including ETS1 and ELK1, were ranked high in normal organoid (K1030N1_GN1) and *VHL* organoids (K1013T2_GB1) (**Figure 4D-E, Table S3**). K1013T2_GB1 is also more similar to normal organoid than other renal cancer organoids in terms of global chromatin landscape (**Figure 4C**). Consistent with observations from scRNA-seq regulon analysis, K1030T2_GB1 (VB) showed higher accessibility in stress response TF JUNB peak regions than K1040T1_GB1 (VP) (**Figure 4F-H**). However, normal organoids also exhibit higher stress response TF activity (**Figure 4B** and **4D**), suggesting that the *PBRM1* mutation might be associated with downregulation of the stress response.

Notably, several renal lineage transcriptional factors, including PAX8, HNF1A and HNF1B, were ranked lower in *BAP1*-driven organoids compared to other genotypes (**Figure 4D-G**). Consistent with this, kidney lineage scores were significantly higher in *PBRM1*-mutated tumourgraft (**Figure S6B**). However, in clinical specimen cohorts, there were no significant differences, likely due to the purity issue. It has been shown that in ccRCC cells, PAX8 and HIF2A cooperate to regulate *CCND1* expression through a distal enhancer^51^. Our data showed that the PAX8-HIF2A binding site is more accessible in K1040T1_GB1 (VP) than in K1030T2_GB1 (VB), thereby allowing *CCND1* expression in K1040T1_GB1 (VP) (**Figure 4I**). We then reasoned that the differential expression of the lineage factor *PAX8* could explain this regulation. Surprisingly, we found that an *antisense PAX8* RNA was expressed in K1030T2_GB1 (VB), leading to *PAX8* silencing (**Figure S6C**). This suggests that *BAP1*-driven organoids are likely to diverge from the renal lineage and achieve malignancy by alternative pathways.

Collectively, transcriptomic and epigenomic profiling data from preclinical model biobank and clinical specimens suggested distinct intrinsic and extrinsic biological programmes driven by *PBRM1* and *BAP1* mutations. Intrinsically, *BAP1* mutant cells exhibit increased proliferation and a stronger stress response, whereas *PBRM1* mutation is associated with renal lineage. While the hypoxia response difference is unclear in bulk data, scRNA-seq suggests that *PBRM1* mutant cells exhibit a stronger hypoxia signature. Extrinsically, *BAP1* mutation is associated with an inflamed TME and *PBRM1* mutation is associated with a vascularised and fibrotic TME.

### PBRM1 loss defines a permissive state

Organoids, paired with genetic engineering, provide a powerful system for studying driver functions in cancer research^52,53^. We adapted a similar approach to investigate the functional consequences of *PBRM1* loss through CRISPR/Cas9-mediated gene editing in organoids with normal and tumour backgrounds (**Figure 5A**). Mutation status at the target loci was confirmed by Sanger sequencing or amplicon sequencing of the *PBRM1* regions (**Figure S7A-D**). Transcriptomic profiling revealed that the dominant source of transcriptional variation lay in the genetic background, and an additional *PBRM1* mutation introduced a subtle transcriptional shift (**Figure 5B, Table S6**). Gene set enrichment analysis confirmed that *PBRM1*-specific genes observed in preclinical models were also enriched in *PBRM1*-knockout organoids compared with parental normal or tumour organoids (**Figure S7E-F**). In K1030T2_GB1 (VB), *PBRM1* knockout resulted in 1217 upregulated and 918 downregulated genes (LFC > 0.5, padj < 0.05) (**Figure S7G**, **Table S6**). Notably, *LOX* and *TGFBI* were again upregulated and appeared to be *PBRM1*-specific targets (**Figure S7H-I**). This is consistent with their upregulation and prevalence of fibrosis in *PBRM1*-driven tumours (**Figure 3B, Figure S5E-F**).

**Figure 5.**
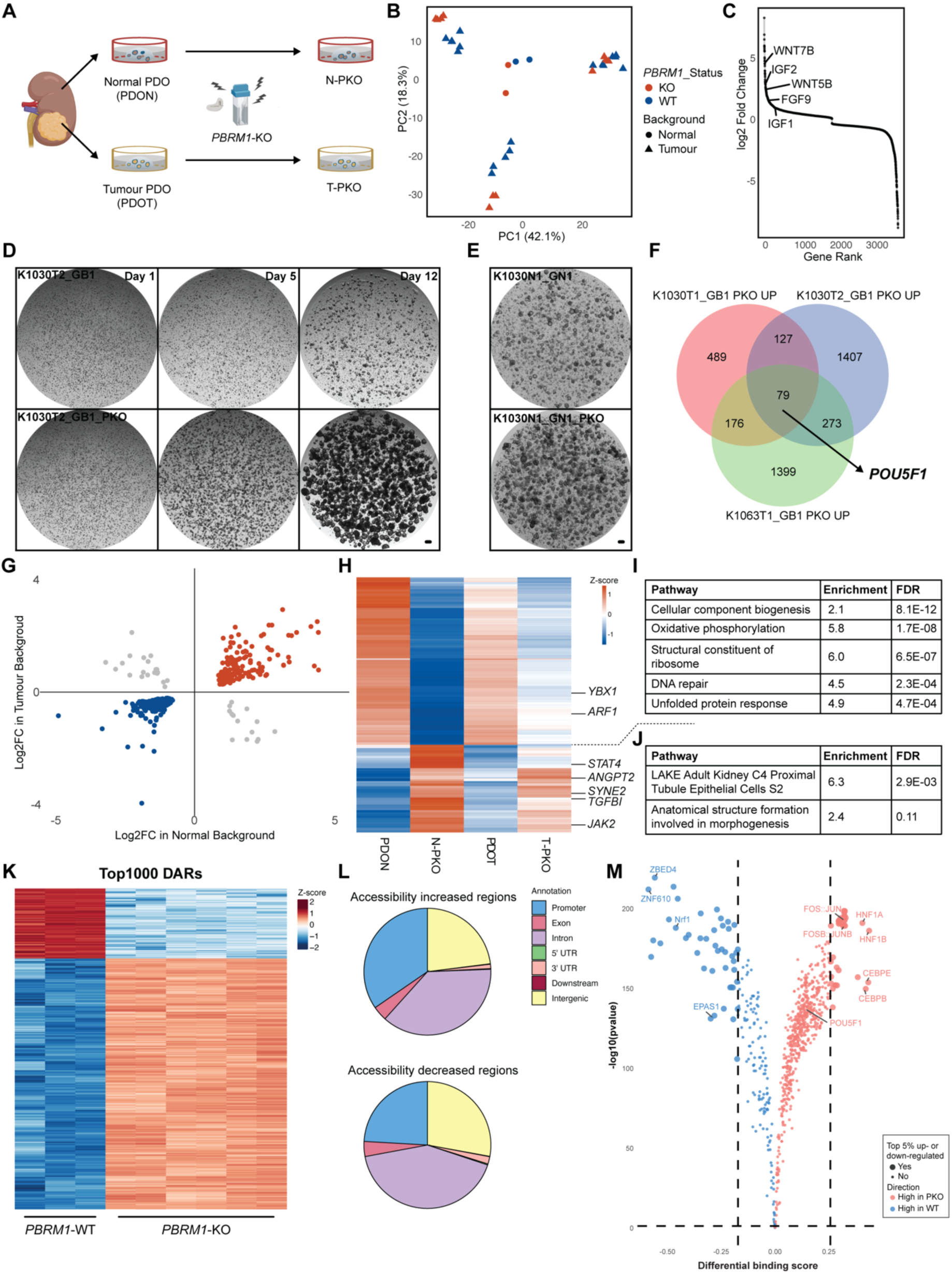
*PBRM1* loss establishes a permissive transcriptional and chromatin state in ccRCC organoids. **(A)** Schematic of CRISPR/Cas9-mediated *PBRM1* knockout (PKO) in patient-derived normal (PDON) and tumour (PDOT) organoids. (B) Principal-component analysis (PCA) of bulk transcriptomes from parental and *PBRM1*-engineered organoid lines. (C) Rank plot showing all differentially expressed genes in *PBRM1*-KO K1030T2_GB1 organoids relative to parental controls. (D and E) Representative bright-field images illustrating enhanced clonogenic potential in *PBRM1*-KO tumour organoids (D) and normal organoids (E). Scale bar, 100 µm. (F) Venn diagram showing shared and unique upregulated genes upon *PBRM1* loss across three independent tumour organoid lines. (G) Scatterplot comparing fold changes of differentially expressed genes upon *PBRM1* loss in tumour and normal organoids.(H) Heatmap of consistently up- and down-regulated genes following *PBRM1* knockout in both backgrounds (sample groups as in A). (I and J) Enriched pathways among consistently upregulated (I) and downregulated (J) genes. FDR, false discovery rate. (K) Heatmap of normalised accessibility scores for the top 1000 differentially accessible regions (DARs) between *PBRM1*-WT and *PBRM1*-KO organoids. (L) Genomic annotation of regions showing increased (top) or decreased (bottom) chromatin accessibility following *PBRM1* loss. UTR, untranslated region. (M) Volcano plot showing differential transcription factor binding activities between *PBRM1*-WT and *PBRM1*-KO organoids. Significance was defined by −log10(p-value) above the 95% quantile and differential binding scores in the top 5% in each direction. See also Figure S7 and Table S6.

Intriguingly, genes involved in growth and stemness signalling, such as *IGF2*, *FGF9*, and WNT ligands, were upregulated upon *PBRM1* loss (**Figure 5C**, **Figure S7G**). Indeed, the *PBRM1*-null engineered cells demonstrated increased organoid recovery from single-cell seeding relative to their parental line, regardless of normal or tumour background, supporting a stemness-associated phenotype (**Figure 5D-E**). The pluripotency marker *POU5F1* (Oct-4) was upregulated upon *PBRM1* loss in all tumour organoids (**Figure 5F**). Consistent with this, single cell regulon analysis also showed POU5F1 as an active TF in VP models (**Figure 4B**). This provides a link between *PBRM1* loss and the stemness programme in renal organoids. Compared to the tumour background, *PBRM1* loss in the normal kidney organoid context led to fewer differentially expressed genes, with a predominance of downregulated over upregulated genes (**Table S6**). We grouped the upregulated and downregulated genes with the same direction in normal or tumour contexts, which revealed 161 co-upregulated genes and 307 co-downregulated genes (**Figure 5G-H**). *JAK2*, *STAT4* and *TGBFI* were among the co-upregulated genes, and *YBX1* was consistently downregulated. Gene set enrichment analysis revealed that downregulated genes were enriched in the processes of protein biogenesis, oxidative phosphorylation, and DNA repair. In contrast, the upregulated genes were enriched in the proximal tubule lineage and morphogenesis process (**Figure 5H-J**).

To gain insights into the regulatory landscape remodelling upon *PBRM1* loss, we performed ATAC-seq to examine the chromatin accessibility changes. Loss of *PBRM1* leads to a significant perturbation to chromatin accessibility, and there is more accessible chromatin compared to *PBRM1* wildtype (**Figure 5K, Table S6**). Moreover, a higher proportion of peak regions with increased accessibility is annotated as promoters (**Figure 5L, Table S6**), which should be associated with gene activation. Differential TF binding analysis revealed HNF1A and HNF1B as the top TFs occupying the *PBRM1* loss-associated peaks (**Figure 5M**). POU5F1 also has a higher binding score in the *PBRM1*-KO organoid, although it is not among the top ones. Together, these findings suggest that upon loss of *PBRM1*, the renal epithelial progenitor property is reinforced. Protein translation is reduced to conserve resources and energy, which should create a permissive state for the pervasive selection and tolerance of multiple subclones during further evolution.

### BAP1 loss drives a progressive phenotype

In contrast to *PBRM1*-knockout cells, which enriched themselves and dominated the population within 20 days, cells with *BAP1* knockout were depleted over time (**Figure 6A-B**). However, this fitness disadvantage can be rescued when knocking out *BAP1* in a tumour background or on the background of *CDKN2A* loss and *VHL* loss (**Figure 6C-D**, **Figure S8A-B**). Organoids with *BAP1* loss also exhibited enrichment of the *BAP1*-specific gene set as identified in *BAP1*-driven organoids (**Figure S8C-D**). Surprisingly, although *BAP1* mutant cells are selected in tumour background, the E2F target gene set is downregulated (**Figure 6E**). This contradicts clinical observations and previous data showing that *BAP1*-driven tumour cells have a higher proliferative capacity. We then examined potential explanations and observed a striking downregulation of genes critical for chromosome segregation, including *CDC20*, *CENPE*, and *CENPF* (**Figure 6F-G, Table S7**). This demonstrates that *BAP1* plays a key role in maintaining chromosomal stability, a prerequisite for proper cell division^54^. The presence of micronuclei, a consequence of chromosomal missegregation, also supports the notion that *BAP1*-mutated organoids are more prone to generate copy number alterations (**Figure 6H**). Moreover, in TRACERx Renal samples with multiregional sampling, we observed a trend that tumours with clonal *BAP1* mutations have a higher number of clonal CNVs (**Figure 6I**, excluding Chr 3-related CNV, p=0.0703), indicating clonal expansion upon *BAP1* loss is coupled with CNV fixation. However, karyotyping of these *BAP1*-knockout lines did not reveal additional chromosomal aberrations (data not shown), indicating that transcriptional priming for chromosomal instability may precede observable genomic alterations.

**Figure 6.**
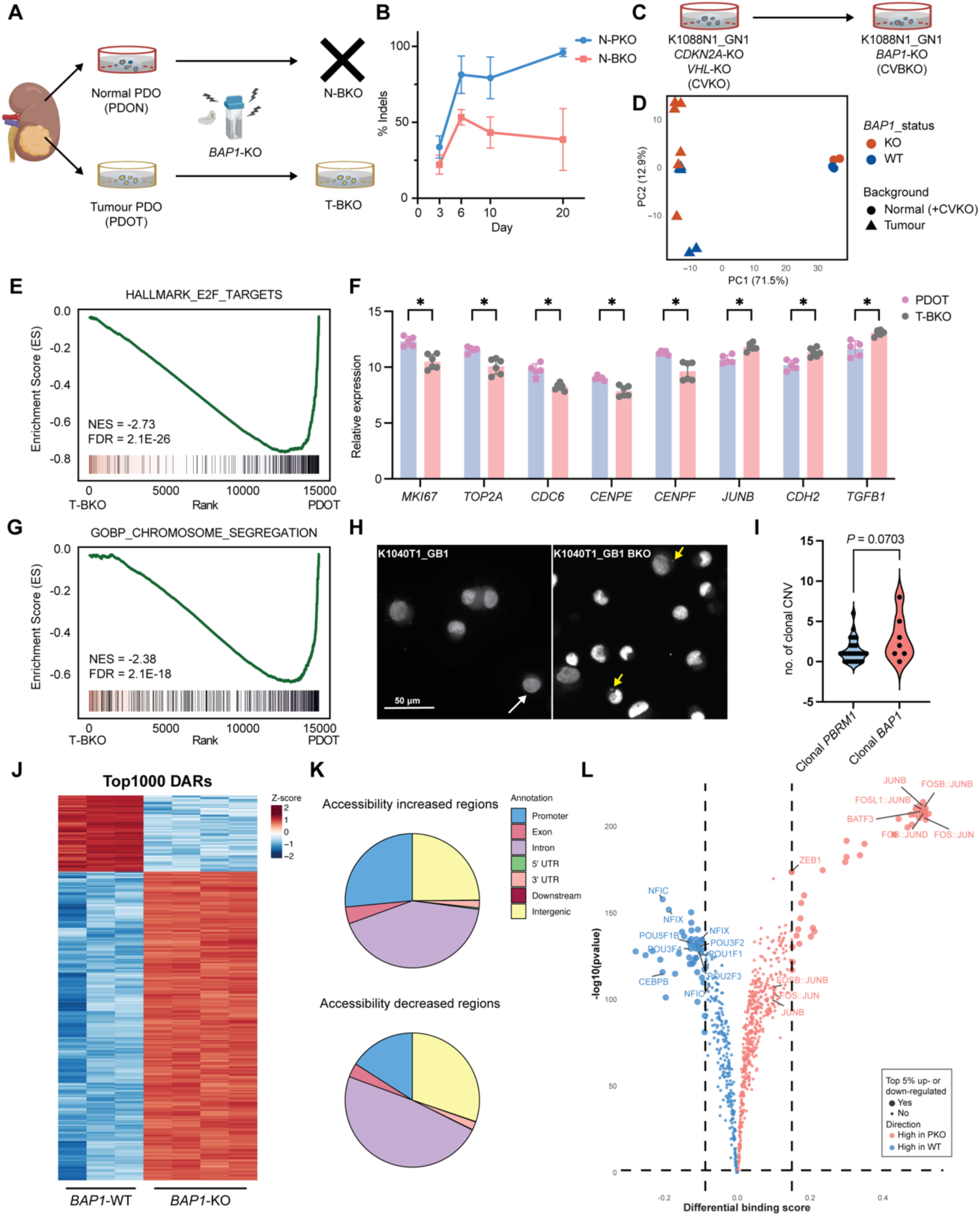
*BAP1* loss drives a progressive phenotype. **(A)** Schematic of CRISPR/Cas9-mediated *BAP1* knockout (BKO) in patient-derived normal (PDON) and tumour (PDOT) organoids. The cross indicates unsuccessful expansion. (B) Tracking of variant allelic frequency (VAF) in normal kidney organoids following electroporation with guide RNAs targeting *PBRM1* or *BAP1*. (C) Schematic of CRISPR/Cas9-mediated *BAP1* knockout (BKO) in normal organoids (K1088N1_GN1) with concurrent *CDKN2A* and *VHL* loss (CV background). (D) Principal-component analysis (PCA) of bulk transcriptomes from parental and *BAP1*-engineered organoid lines. (E) GSEA of *BAP1*-KO tumour organoids versus parental controls using the Hallmark E2F Targets gene set. (F) Comparison of normalised expression (DESeq2 VST) of selected genes between *BAP1*-KO and parental tumour organoids (*n* = 5 PDOT, *n* = 6 PDOT-BKO). Unpaired *t*-test; *: *p* < 0.01. (G) GSEA of *BAP1*-KO tumour organoids versus parental controls using the Gene Ontology Biological Process (GOBP) Chromosome Segregation gene set. (H) Representative DAPI staining showing micronucleus presence in *BAP1*-KO and parental tumour organoids. Scale bar, 50 µm. (I) Violin plot showing the number of clonal copy-number variants (CNVs), excluding chromosome 3p loss, in tumours with clonal *BAP1* or *PBRM1* mutations. Unpaired *t*-test; *p* = 0.0703. Clonal *PBRM1*, *n* = 25; clonal *BAP1*, *n* = 7. (J) Heatmap showing normalised accessibility scores for the top 1000 differentially accessible regions (DARs) between *BAP1*-WT and *BAP1*-KO organoids. (K) Genomic annotation of regions with increased (top) or decreased (bottom) chromatin accessibility following *BAP1* loss. UTR, untranslated region. (L) Volcano plot showing differential transcription-factor binding activities between *BAP1*-WT and *BAP1*-KO tumour organoids. Significance was defined by −log10(p-value) above the 95% quantile and differential binding scores in top 5% in each direction. See also Figure S8 and Table S7.

Notably, the mesenchymal marker *CDH2* (encoding N-cadherin) was consistently upregulated, along with *TGFB1* and genes associated with TGF-β signalling (**Figure 6F**, **Figure S8F**). This indicates *BAP1* loss may reprogram cells towards EMT. ATAC-seq on paired *BAP1*-WT and *BAP1*-KO tumour organoids revealed a more accessible chromatin landscape (**Figure 6J-K, Table S7**). Importantly, genes near accessible peaks are enriched in the TGF-β signalling pathway (**Figure S8H**). This supports a direct regulation of TGF-β signalling by the *BAP1* gene, and loss of *BAP1* promotes a mesenchymal and invasive phenotype. Furthermore, this also explains the presence of high TGF-β activity LRRC15+ myofibroblasts in *BAP1* tumours (**Figure 3H**).

Lastly, the stress response indicator *JUNB* was also upregulated upon *BAP1* loss (**Figure 6F**), consistent with previously observed transcriptional profiles of *BAP1*-driven tumours. In line with this, transcriptional factor units of activator protein-1 (AP-1), the Fos/Jun cluster, were shown to have a significantly higher binding score in *BAP1*-KO organoids compared to WT organoids (**Figure 6L, Table S7**). Moreover, the IL2-STAT5 signalling pathway is enriched in *BAP1*-KO organoids (**Figure S8E**). The AP-1 (Fos/Jun) has been shown to regulate cytokine expression and modulate inflammatory response^55,56^. It has also been shown that *BAP1*-tumours are more inflamed^57^ (**Figure 3B and 3F**). Our data provide a functional link between *BAP1* loss and an inflammatory phenotype via tumour-intrinsic AP-1 activation.

## DISCUSSION

The transition from an indolent stage to aggressive ccRCC represents a pivotal stage in tumour evolution, driven by the acquisition of additional somatic alterations that confer selective advantages and shape disease phenotype. However, functional investigation into these genotype-phenotype relationships has been limited, partly due to the lack of physiologically relevant experimental models, which has consequently hampered the development of tumour-intrinsic targeted therapies. Here, we build a preclinical model biobank for ccRCC studies and systematically demonstrate the functional consequences of *PBRM1* and *BAP1* loss by integrating genomic and transcriptomic data with functional perturbations in preclinical models.

Our data, together with the established consensus^16,58^, support the existence of two distinct evolutionary trajectories in ccRCC, shaped by *PBRM1* and *BAP1* mutations. *PBRM1*-driven tumours behave as passive responders, maintaining and modestly modulating the transformed phenotype initially established by *VHL* loss. This trajectory is associated with a small but sustained fitness advantage fuelled by a supportive fibrotic niche and epithelial stemness (**Figure 7**). This enables long-term persistence and gradual evolution. Consistent with this, *PBRM1* mutations are observed in approximately 40% of ccRCC cases, representing a prevalent and pervasive evolutionary route.

**Figure 7.**
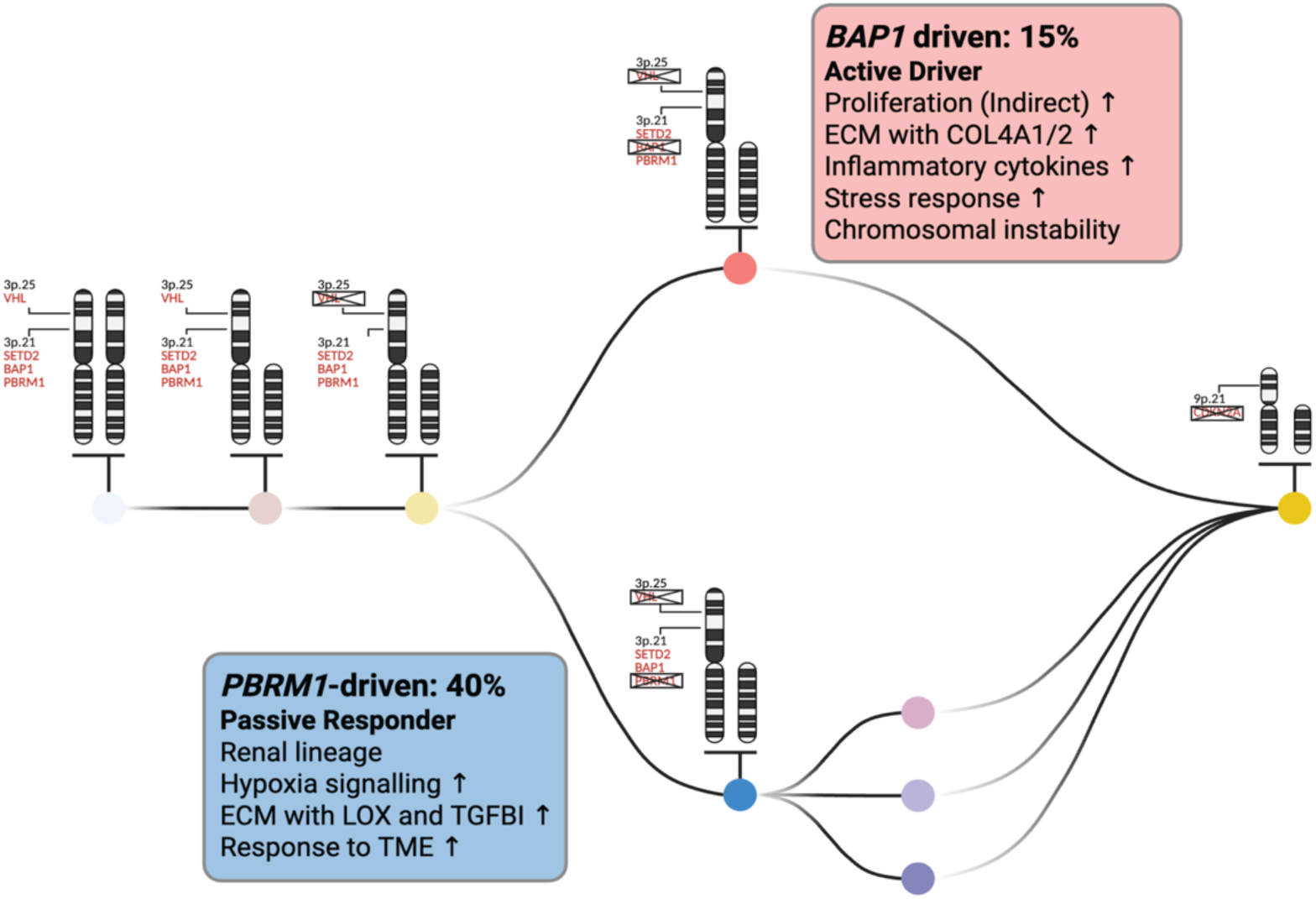
Phenotypic characteristics of two major ccRCC evolutionary routes. Schematic summary of *BAP1*- and *PBRM1*-specific molecular phenotypes defining distinct evolutionary trajectories in ccRCC.

In contrast, *BAP1*-driven tumours follow an active trajectory marked by rapid and aggressive transformation. While *BAP1* mutations confer phenotypic changes, including lineage reprogramming, stress response activation, and CIN, these changes also impose an initial fitness cost (**Figure 7**). Interestingly, a similar fitness cost was reported in normal liver organoids, where *TP53* deficiency rescued the viability of *BAP1*-knockout liver organoids^59^. These observations suggest that only a subset of *BAP1*-mutant cells may be able to overcome this bottleneck; those that do are primed for fast progression. This explains why *BAP1*-mutant tumours, though less frequent, account for roughly 15% of ccRCC cases, tend to be clinically aggressive and are associated with poor outcomes.

Future work should build upon this framework in several directions. Establishing organoid models that represent early-stage tumours would offer critical insights into the initial steps of tumour evolution and provide valuable resources for targeted genetic engineering to recapitulate evolutionary trajectories. Incorporation of tumour-extrinsic components in functional modelling needs to be invested in to fully understand how *BAP1* and *PBRM1* interact with their environment and explore the therapeutic vulnerability. Recent advances in organoid co-culture systems^60^, as well as patient-derived fragment culture systems^61^, might shed light on this. Moreover, genotype-guided therapy could be explored with the framework we developed. For example, the usage of HIF2α inhibitor^62,63^ and lysyl oxidase inhibitor^64^ for *PBRM1*-mutant tumours could be considered.

## Limitations of the study

Nonetheless, several limitations of this study must be acknowledged. First, although the preclinical model biobank successfully recapitulated the major ccRCC genotypes, most models represented advanced disease stages characterised by extensive CNVs. This inherent bias limits insight into the early events of tumour evolution and may miss the initial phenotypical adaptations after the occurrence of genetic drivers. Second, although preclinical tumour models offer a powerful platform for functional interrogation, they lack immune and stromal components, which are essential to ccRCC biology. Consequently, certain transcriptional features observed in clinical specimens, particularly those related to intercellular signalling and inflammatory pathways, may reflect tumour–microenvironment interactions absent *in vitro*. Nevertheless, the integration of preclinical models and clinical data suggests a model in which *BAP1*-mutant tumours actively shape their microenvironment, whereas *PBRM1*-mutant tumours rely more on it.

## ACKNOWLEDGMENTS

We are grateful to the patients and their families who generously participated in the TRACERx Renal Study. We thank the Turajlic Lab for fruitful discussion and valuable feedback. We thank Dr Illaria Malanchi and Dr Greg Findlay from the Francis Crick Institute, London, UK, Dr Chris Tape from UCL Cancer Institute, London, UK, as well as Dr Wei Chen, Dr Liang Fang, Dr Qionghua Zhu, Zhiyuan Sun and Yanping Li from SUSTech, Shenzhen, China, for insightful discussions. We thank Andrew Porter from The CRUK Manchester Institute, Manchester, UK, for his helpful feedback on the manuscript. We thank the support from the Science Technology Platforms at the Francis Crick Institute, London, UK, especially the Genomics Facility, the Flow Cytometry Facility, and the Experimental Histopathology Facility. BioRender icons were used in figure preparation under the Francis Crick Institute’s subscription license. This work was supported by the Francis Crick Institute, which receives its core funding from Cancer Research UK, the UK Medical Research Council and the Wellcome Trust (CC2044, S.T.), and by the NIHR BRC (Research project B217, D.D. and S.T.).

## AUTHOR CONTRIBUTIONS

Conceptualization, S.T. and D.D.; methodology, D. D., H. F., A. F., Y. D., F. B., C. S., A. M., A. F. S., and S.T.; Investigation, D. D., H. F., A. F., O. B., S. T. C. S., H. F., A. C., H. P., I. L., K. M., Q. G., M. E., J. W., A. F., S. B. A., Z. T., and B. I. Y.; resources, T. B., S. H., I. S, K. E., E. C., A. M., J. K., A. S. S., L. B., J. L., S. R., A. B., and S. T.; data curation: D. D., H. F., C. S., A. M., A. F. S., A. C., J. W., and I. S.; writing—original draft, D.D. and S.T.; writing—review & editing, D.D., S.T., and H. F. with contributions from all the authors.

## SUPPLEMENTAL INFORMATION

Table S1. Patient and model information.

Table S2. PDO single cell metadata and regulon scores.

Table S3. Transcriptomic and epigenetic characterisation of PDO biobank.

Table S4. Metadata and ssGSEA scores of TCGA, TRACERx Renal, and Tumourgraft cohorts.

Table S5. Curation of public single cell data and differentially expressed genes.

Table S6. Comparative analysis of *PBRM1*-KO and parental organoids.

Table S7. Comparative analysis of *BAP1*-KO and parental organoids.

Table S8. Medium formula and oligo sequences.

---

**Figure S1.**
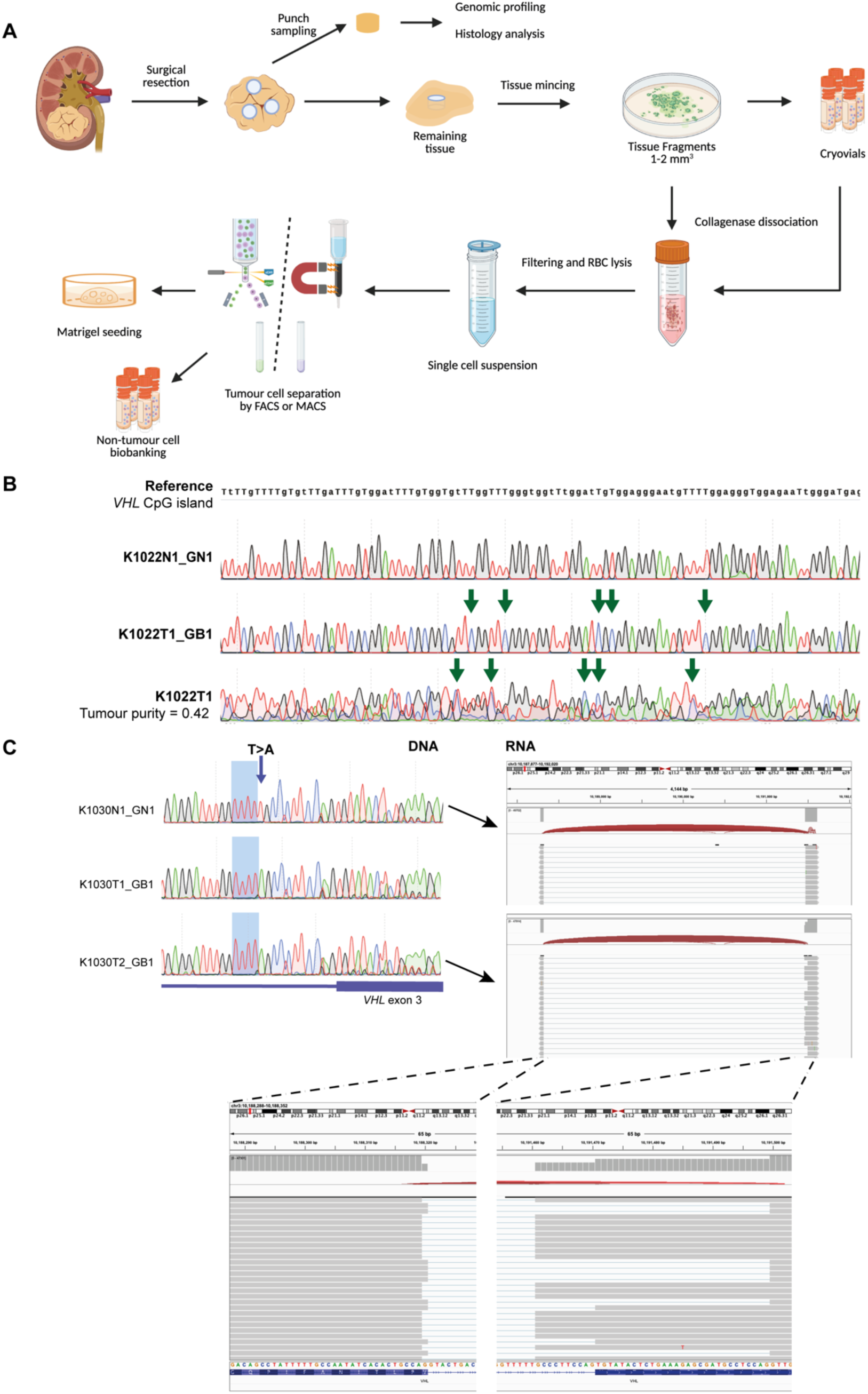
*VHL* inactivation in PDOs. **(A)** Schematic showing experimental procedures of preclinical model derivation. (B) Sanger sequencing traces showing *VHL* promoter methylation detected in the K1022 primary tumour and corresponding tumour-derived organoid (K1022T1_GB1) but absent in the matched normal organoid (K1022N1_GN1). (C) Sanger sequencing traces and Integrative Genomics Viewer (IGV) plots demonstrating an intronic *VHL* mutation that disrupts normal splicing between exons 2 and 3.

**Figure S2.**
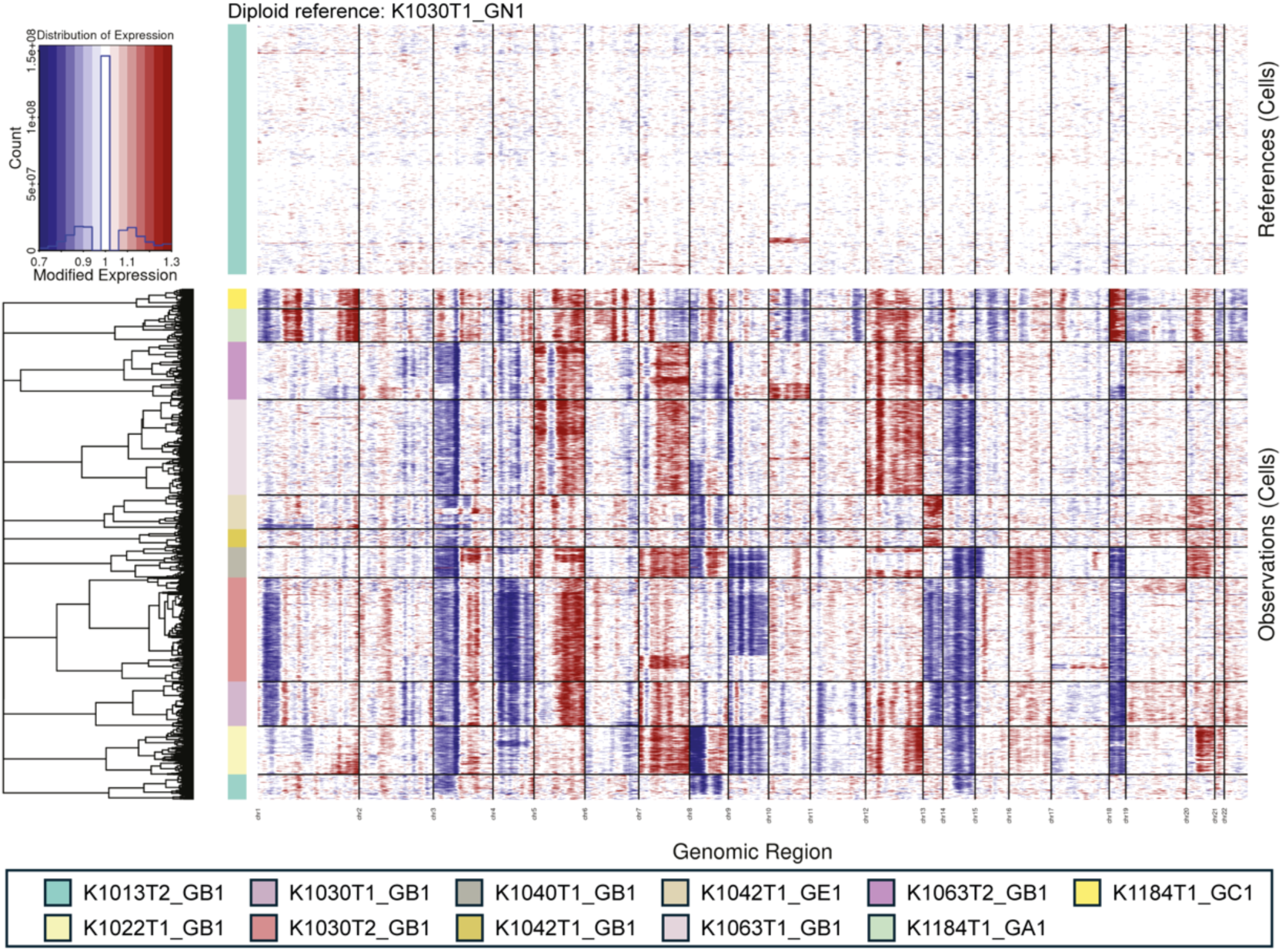
Inferred copy number profiles from single-cell transcriptomes of PDOs. Heatmap showing inferred single-cell copy number variation (CNV) profiles of PDOs using inferCNV, with the normal organoid K1030N1_GN1 used as the diploid reference.

**Figure S3.**
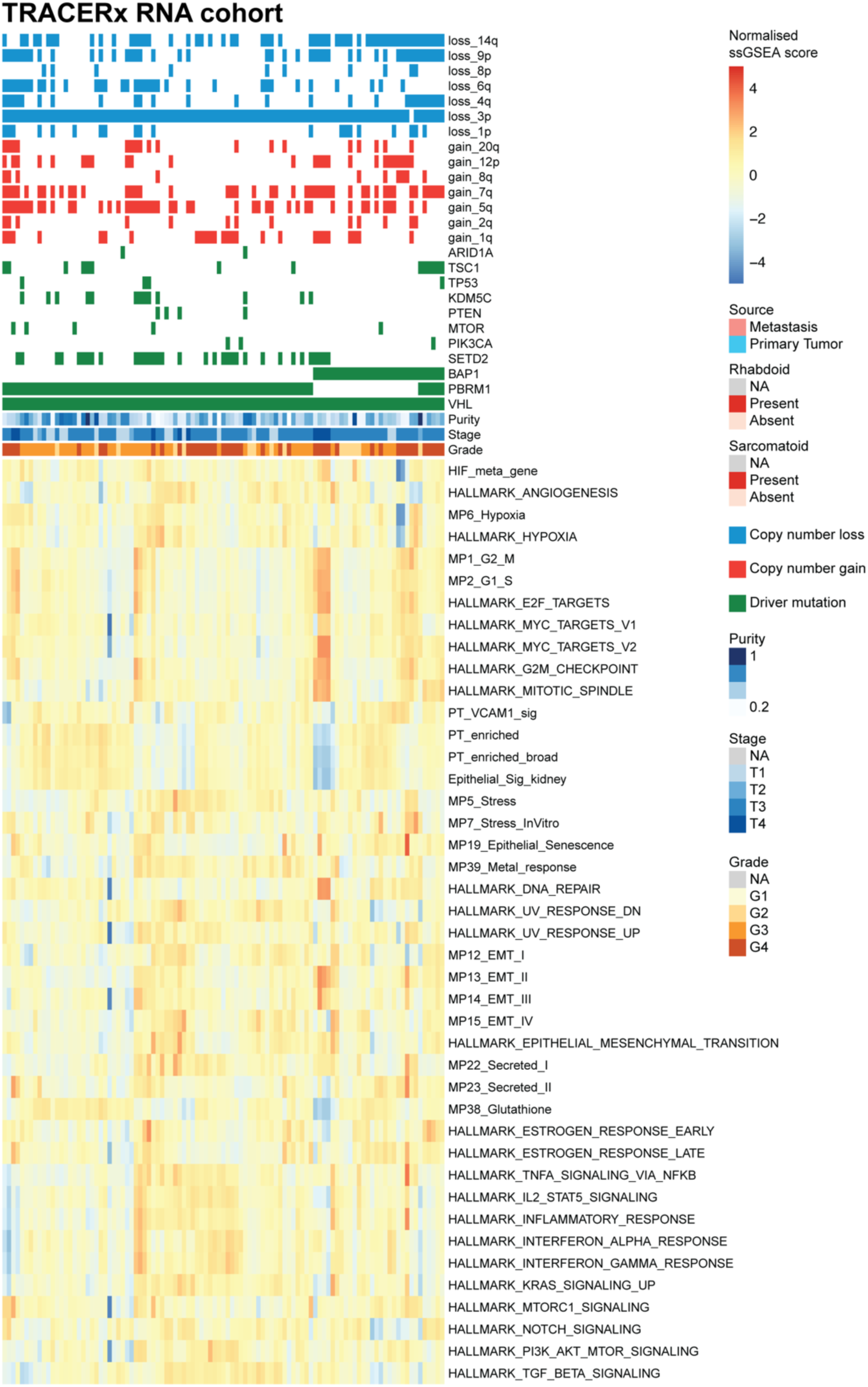
Transcriptional programme landscape in TRACERx RNA cohort. Heatmaps showing normalised single-sample Gene Set Enrichment Analysis (ssGSEA) scores of curated gene sets across clinical samples in the TRACERx Renal cohort. Sample annotations and mutation status are indicated above the heatmaps.

**Figure S4.**
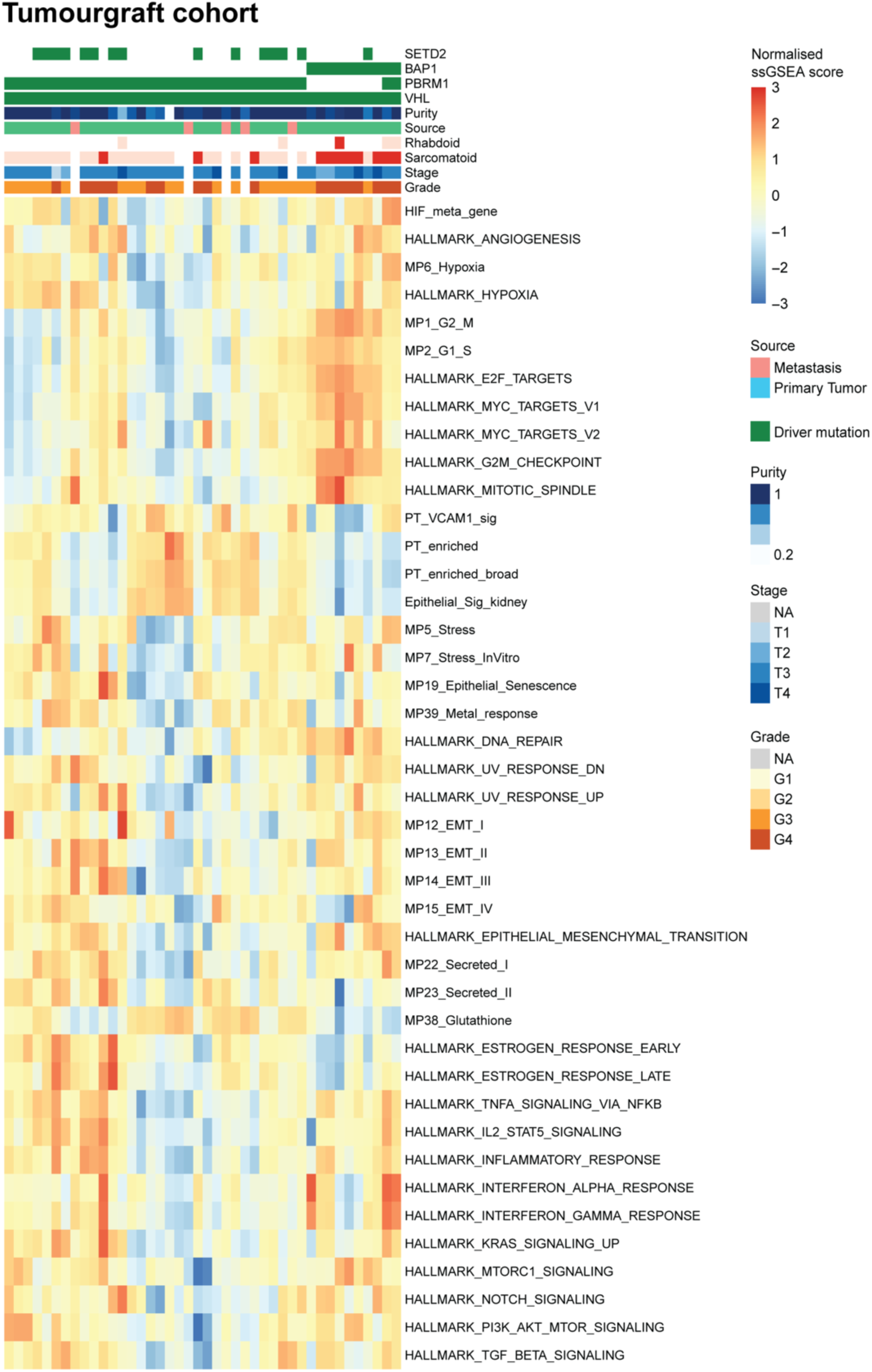
Transcriptional programme landscape in Tumourgraft cohort. Heatmaps showing normalised single-sample Gene Set Enrichment Analysis (ssGSEA) scores of curated gene sets across clinical samples in the Tumourgraft cohort. Sample annotations and mutation status are indicated above the heatmaps.

**Figure S5.**
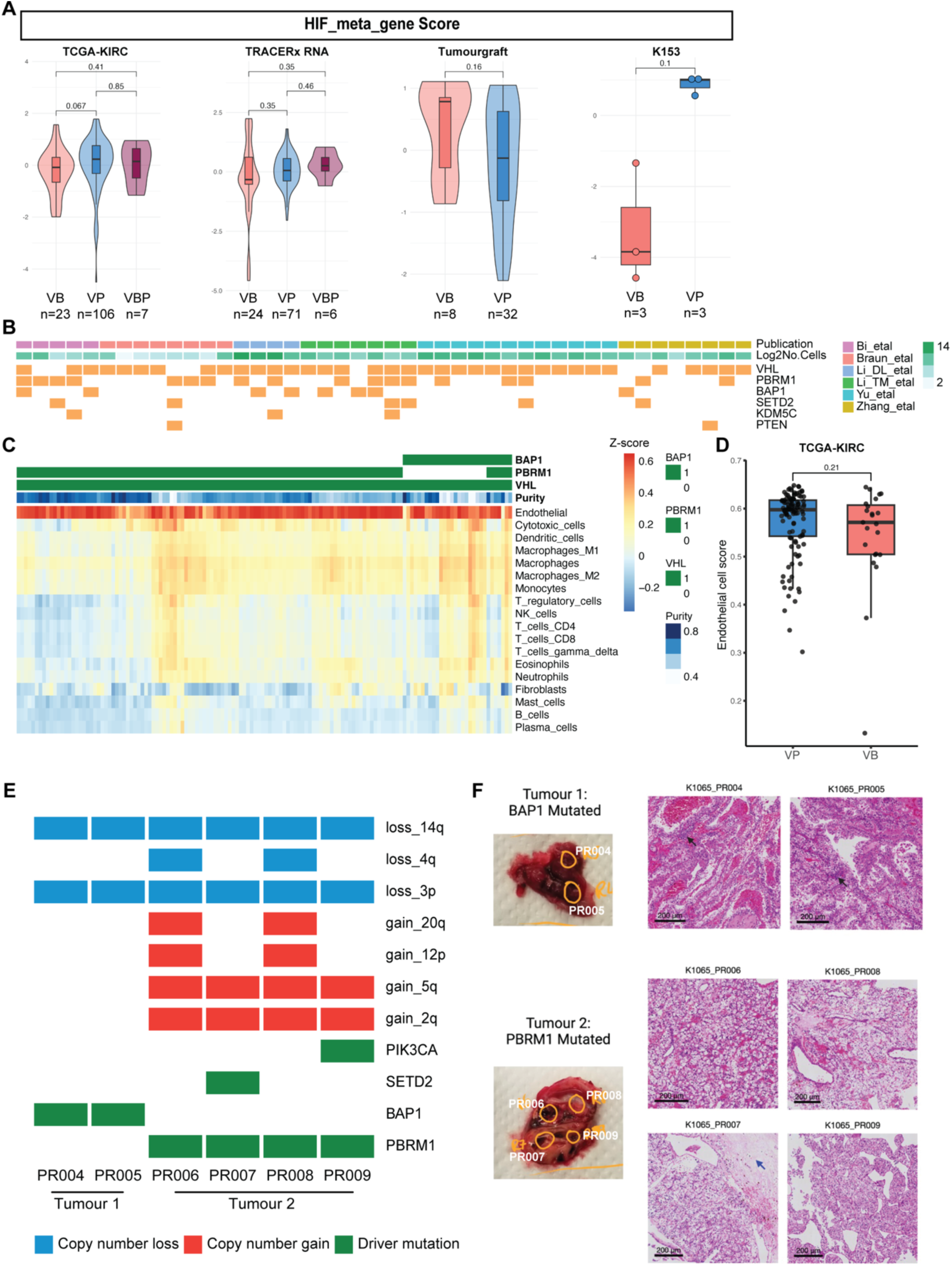
*BAP1*- and *PBRM1*-specific tumour microenvironments. (A) Comparison of hypoxia response (HIF metagene score) between *BAP1*-driven and *PBRM1*-driven tumours in TCGA-KIRC, TRACERx RNA, Tumourgraft cohorts and case K153. Sample sizes are denoted under the group names (B) Summary of the genomic landscape and cell numbers across the integrated scRNA-seq datasets. (C) Heatmap showing the TME composition of *BAP1*-driven and *PBRM1*-driven tumours in TCGA-KIRC. Non-tumour cell population abundance was estimated by ConsensusTME and scaled across tumour samples. (D) Box plot comparing endothelial cell score between *BAP1*-driven and *PBRM1*-driven tumours in TCGA-KIRC cohort (VB: n=23, VP: n=106, Wilcoxon signed-rank test, p=0.21). (E) Oncoplot showing mutation profiles of two separate tumours from VHL patient case K1065 (*BAP1*-mutant tumours: PR004, PR005; *PBRM1*-mutant tumours: PR006, PR007, PR008, PR009). (F) Representative H&E staining showing features associated with *PBRM1*-mutant tumours (clear cells, compact small nests, sclerotic stroma; blue arrows) and *BAP1*-mutant tumours (granular cytoplasm, papillary architecture, increased immune infiltrates; black arrows) in case K1065.

**Figure S6.**
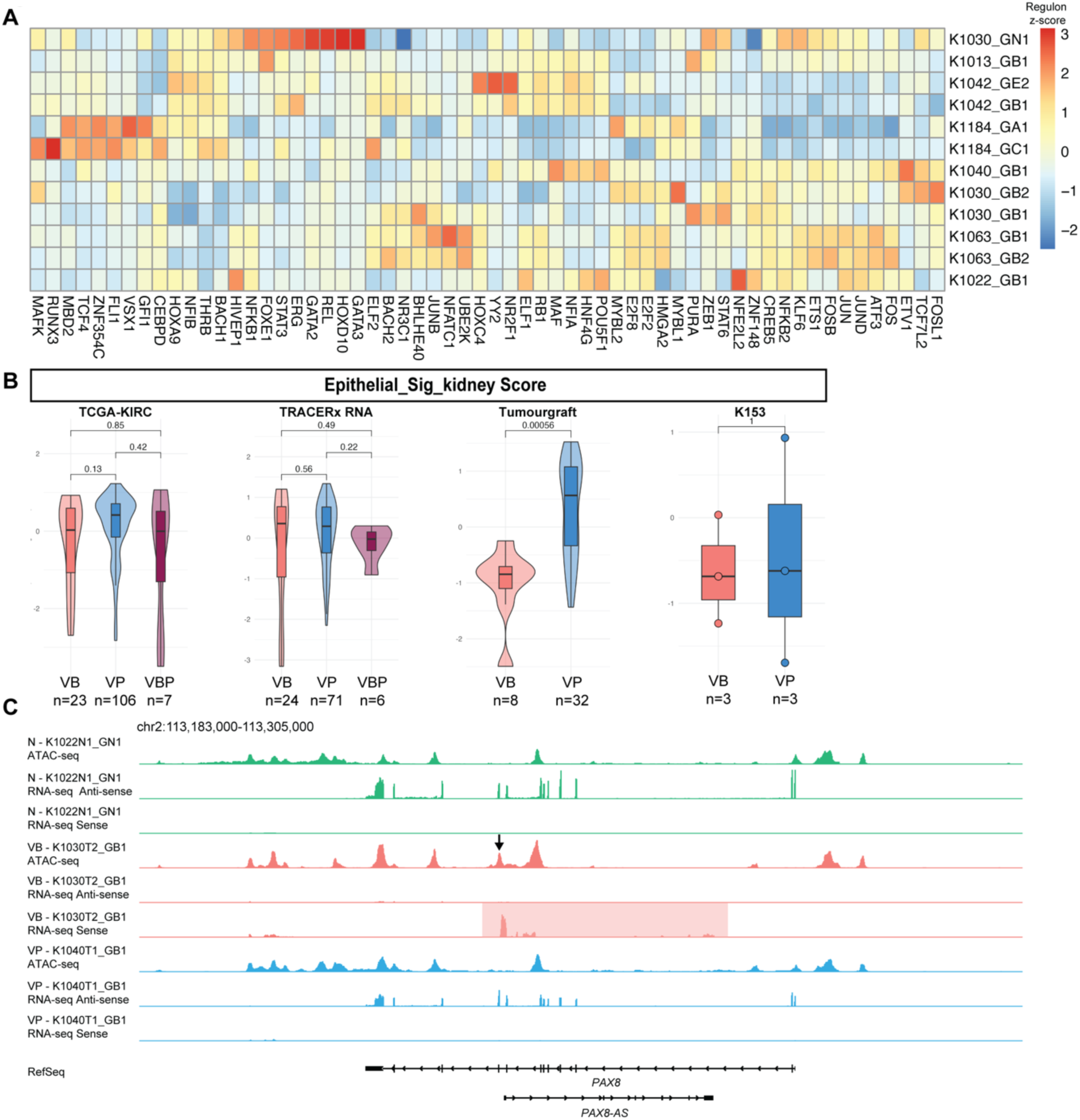
Renal lineage factor activity in *BAP1*- and *PBRM1*-mutant cells. (A) Heatmap showing the *z*-scores of the top 60 variable regulons across preclinical model lines. (B) Comparison of renal lineage programme score (kidney epithelial signature) between *BAP1*-driven and *PBRM1*-driven tumours in TCGA-KIRC, TRACERx RNA, Tumourgraft cohorts and case K153. Sample sizes are denoted under the group names. (C) Coverage plots displaying ATAC-seq chromatin accessibility and RNA-seq expression of *PAX8* and *PAX8-AS* across normal (N), *BAP1*-driven (VB), and *PBRM1*-driven (VP) organoids.

**Figure S7.**
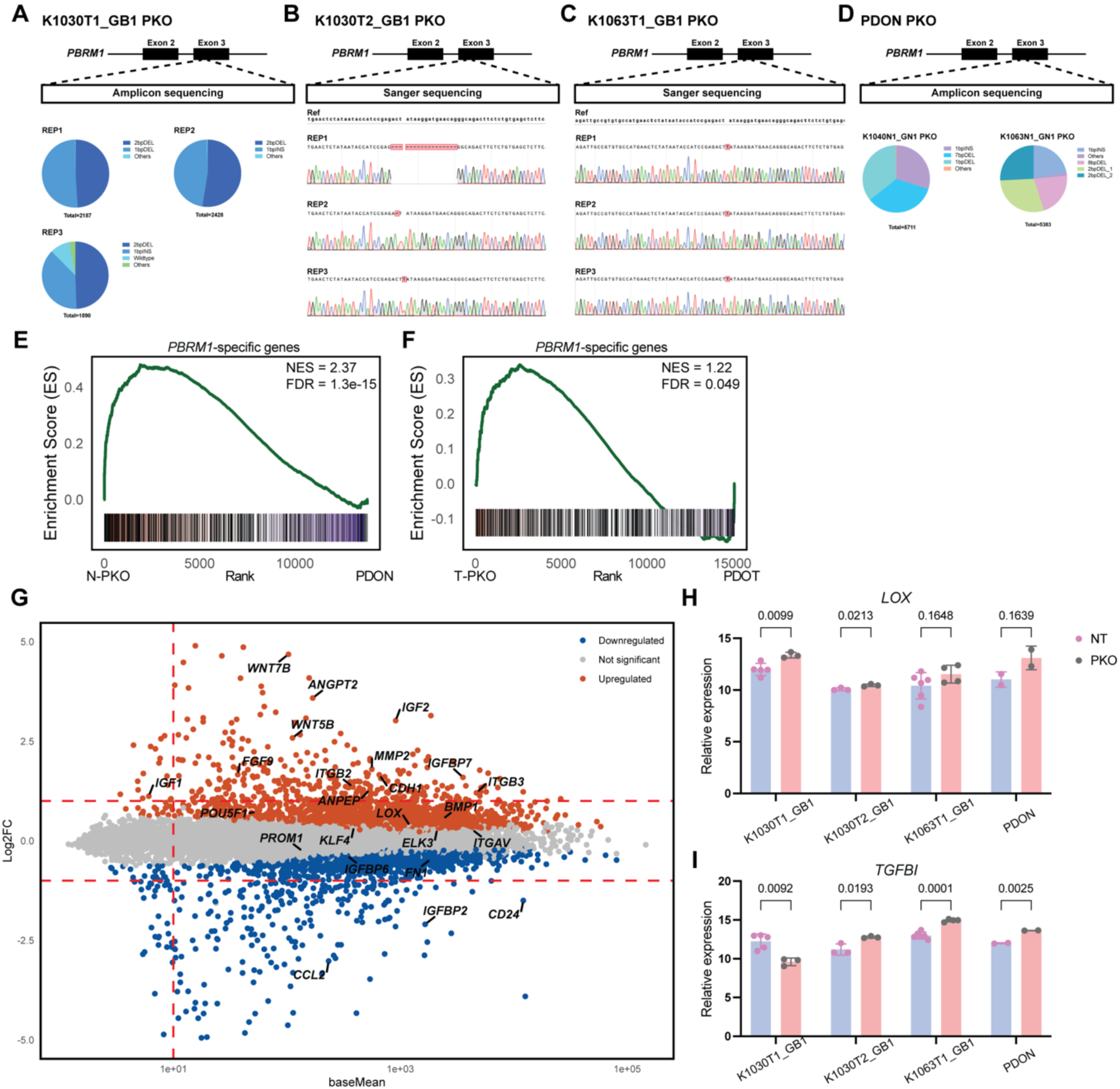
*PBRM1* engineering in organoids. **(A–D)** Variant pie charts and Sanger sequencing traces confirming *PBRM1* knockout in (A) K1030T1_GB1, (B) K1030T2_GB1, (C) K1063T1_GB1 and (D) normal renal organoids. (E-F) GSEA of *PBRM1*-KO (E) normal organoids and (F) tumour organoids versus parental controls using the *PBRM1*-specific genes identified in Figure 2E. (G) Mean–difference (MA) plot showing differentially expressed genes in K1030_GB2 *PBRM1*-KO organoids compared with the parental line. (H–I) Expression levels (DESeq2 VST values) of (H) *LOX* and (I) *TGFBI* in *PBRM1*-KO tumour organoids versus parental organoids. Unpaired *t*-test; *p* values are indicated above. Each data point represents one sample.

**Figure S8.**
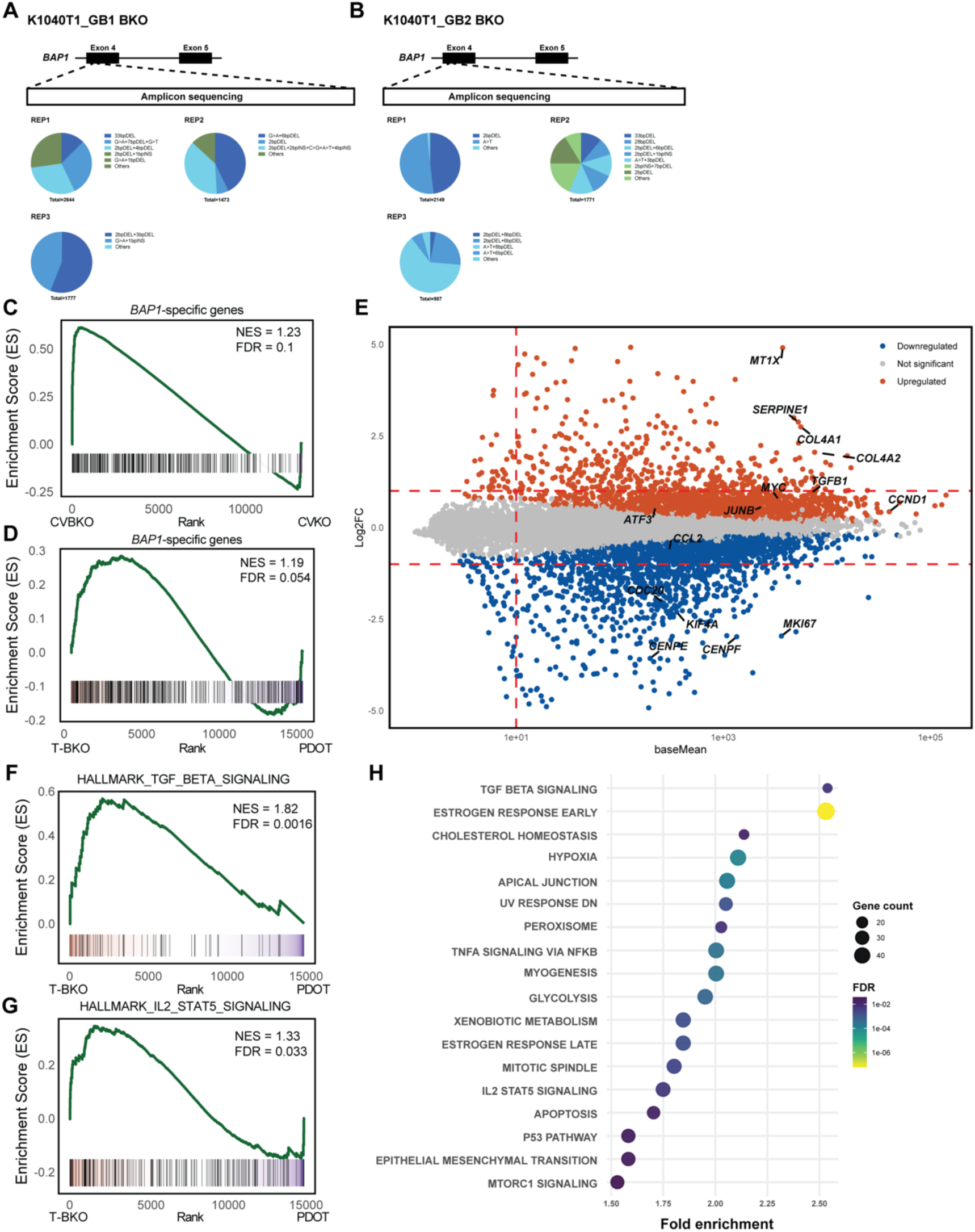
*BAP1* loss activates TGF-β signalling. **(A–B)** Variant pie charts and Sanger sequencing traces confirming *BAP1* knockout in K1040T1_GB1 (A), K1040T1_GB2 (B). (C-D) GSEA of *BAP1*-KO (C) normal organoids and (D) tumour organoids versus parental controls using the *BAP1*-specific genes identified in Figure 2E. (E) Mean–difference (MA) plot showing differentially expressed genes in K1040_GB1 *BAP1*-KO organoids compared with the parental line. (F–G) Gene enrichment analysis of *BAP1*-KO tumour organoids versus parental organoids using the *HALLMARK_TGF_BETA_SIGNALING* (F) and *HALLMARK_IL2_STAT5_SIGNALING* (G) gene sets. (H) Enriched hallmark pathways among genes with increased chromatin accessibility. FDR, false discovery rate.

## STAR★METHODS

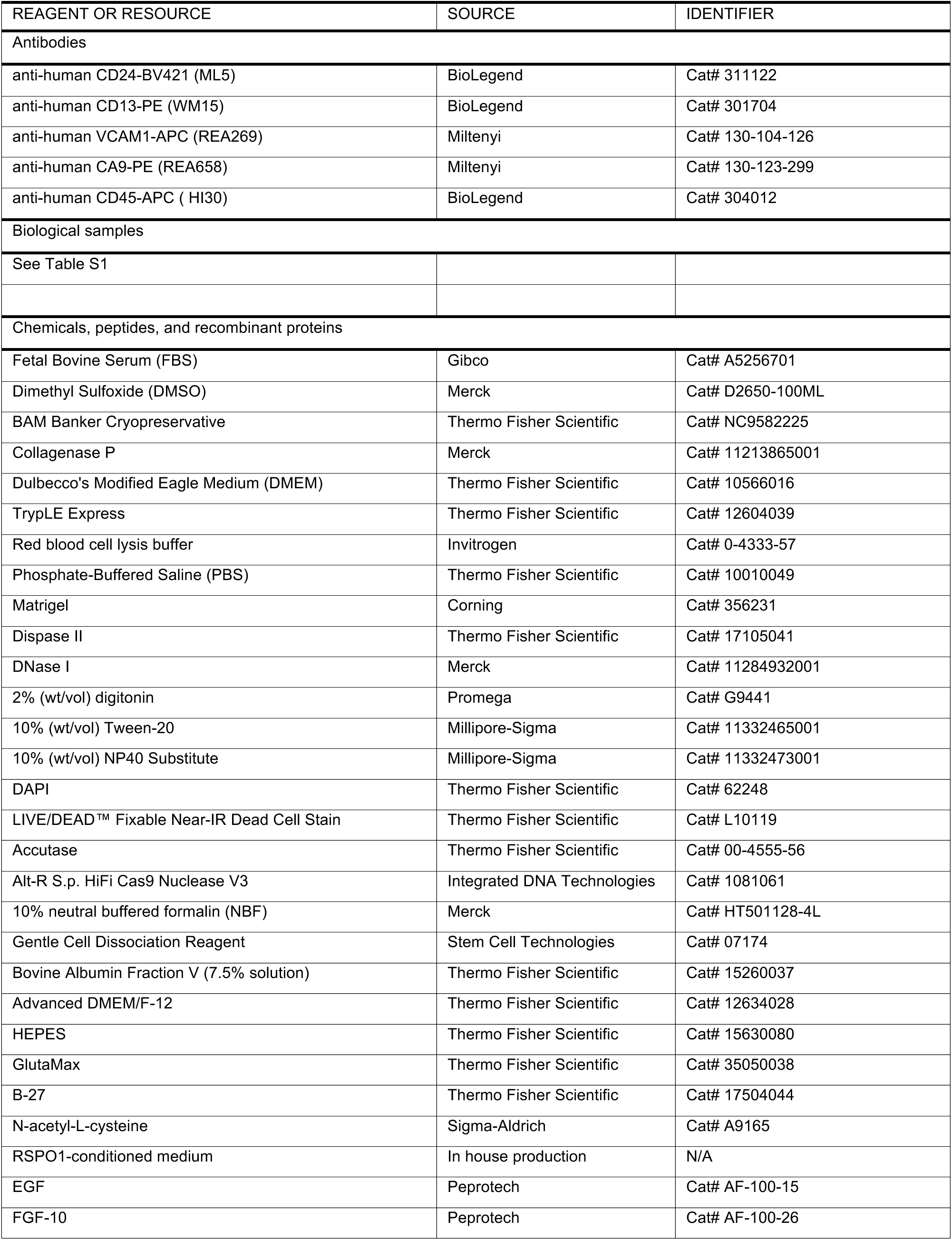
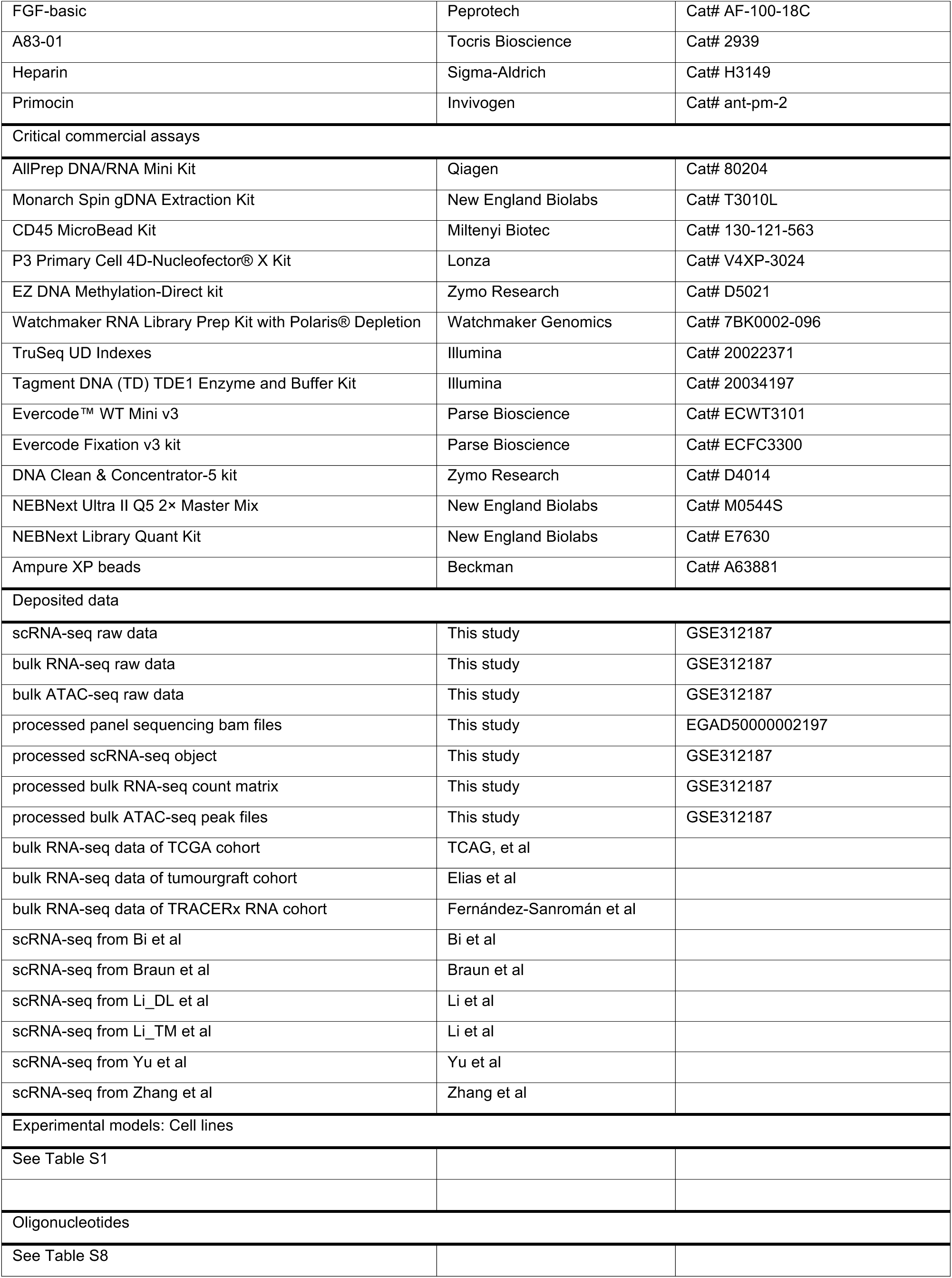
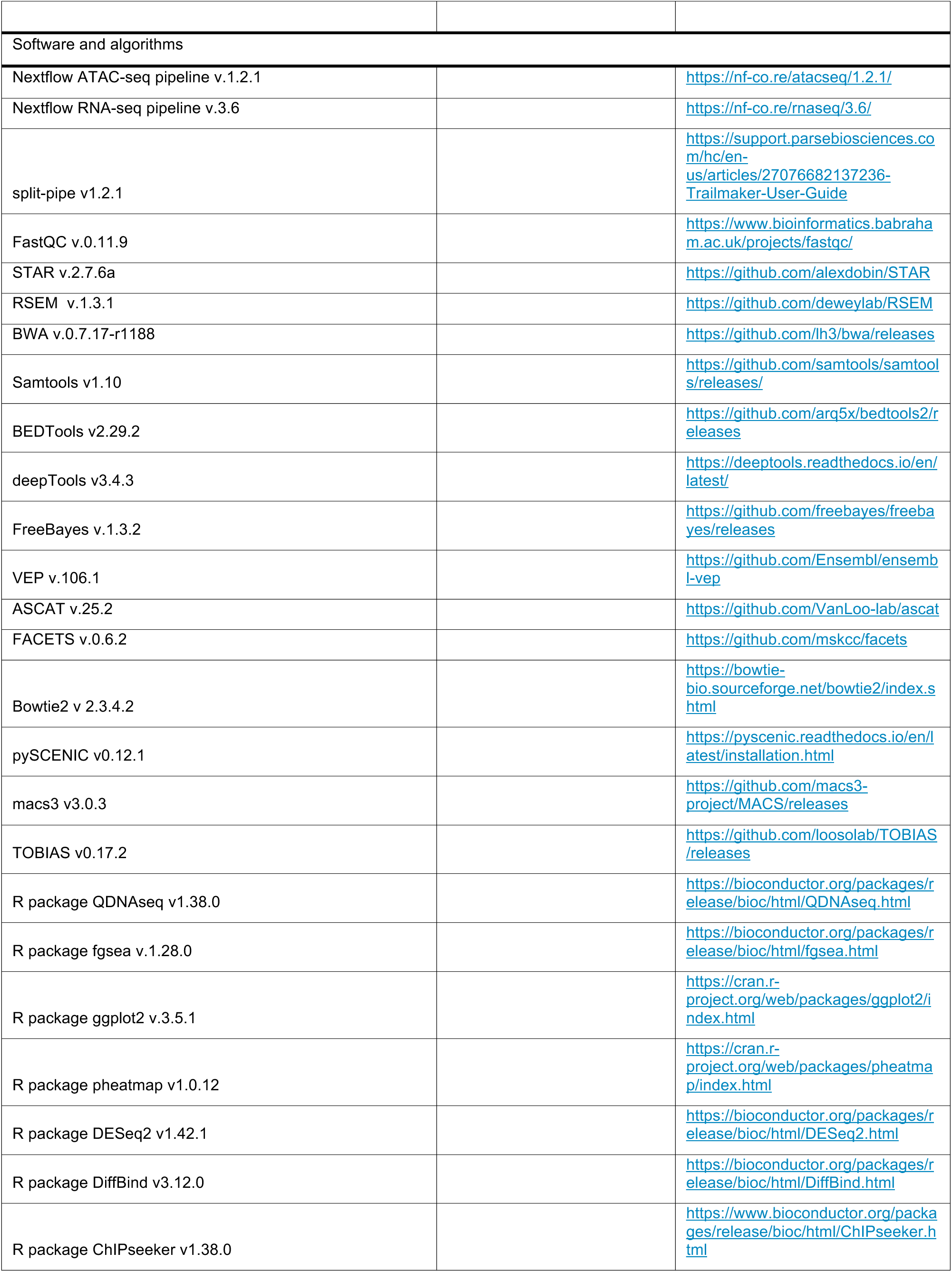
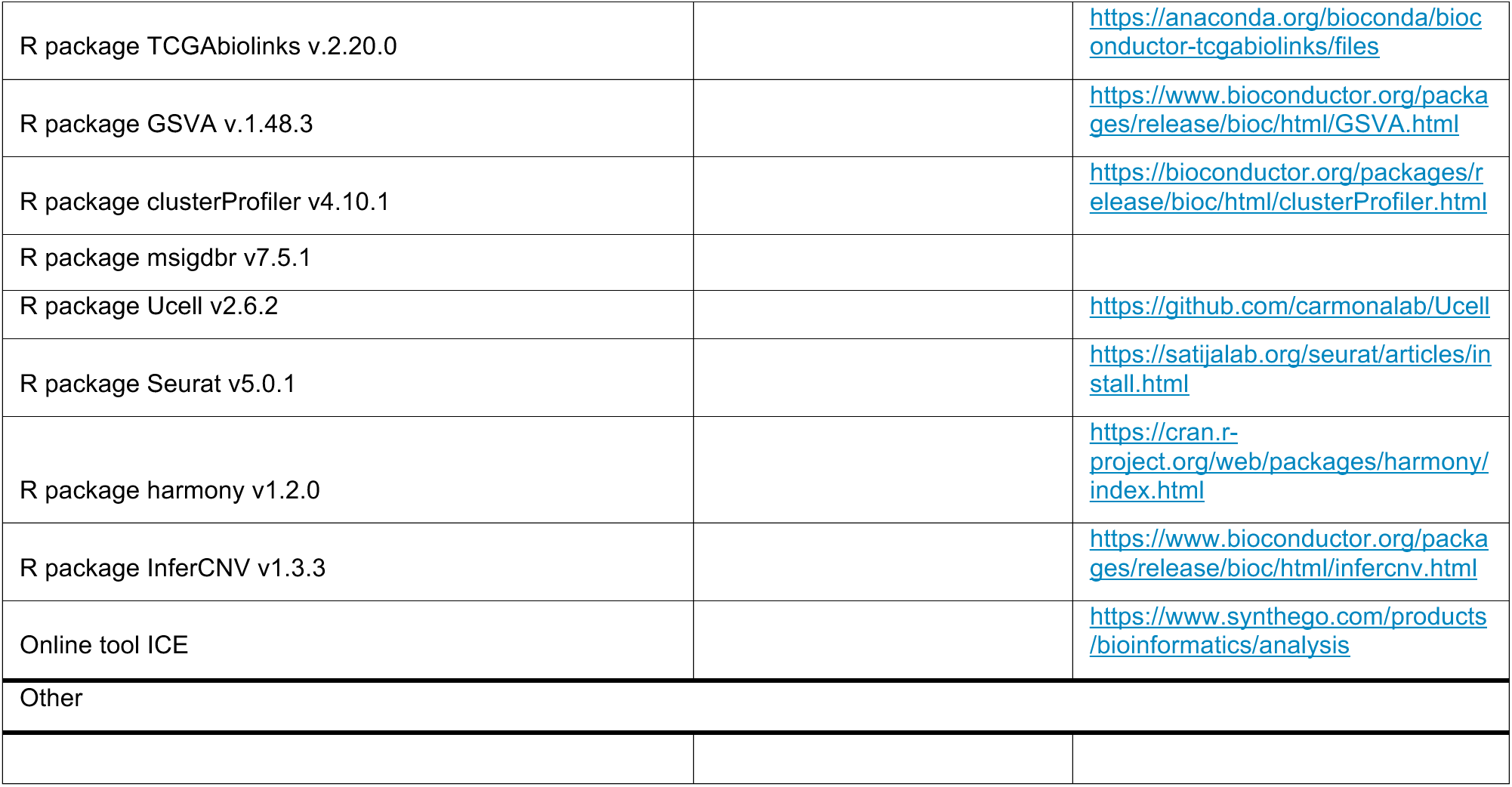

### EXPERIMENTAL MODEL AND STUDY PARTICIPANT DETAILS

#### Clinical Samples

Patients were recruited through the TRACERx Renal study (NCT03226886), an ethically approved prospective cohort study (National Health Service Research Ethics Committee approval 11/LO/1996) sponsored by the Royal Marsden NHS Foundation Trust. Patients were recruited at Royal Marsden Hospital NHS Foundation Trust (RMH), Guy’s and St Thomas’ Hospital NHS Foundation Trust (GSTT), and Royal Free Hospital NHS Foundation Trust (RFH). Every patient in the study was assigned a unique study identity number (ID), with a prefix (“K”) followed by four numbers. All related biological materials, including patient-derived tumour fragments (PDTFs), organoids, and cell lines, were labelled with the corresponding study ID to ensure systematic integration within a patient-centred database.

### METHOD DETAILS

#### Surgical Specimen Processing

Fresh specimens after surgery were examined by a pathologist and sectioned at the hospital sites. One tumour slice was selected for sampling, from which multiple 6-mm punch biopsies were obtained. These biopsies were then vertically halved, with one portion snap-frozen and the other fixed in formalin. The surrounding tissue from the punch biopsy sites, referred to as “donut samples,” was excised and further minced into 1–2 mm³ fragments to generate PDTFs as previously described^66,67^. When available, adjacent normal tissues were also collected and processed in a similar manner. PDTFs were either used immediately for downstream applications or cryopreserved in cryovials containing freezing medium (90% fetal bovine serum (FBS) and 10% dimethyl sulfoxide (DMSO)). Cryopreserved PDTFs were stored in liquid nitrogen for long-term preservation until thawed for dissociation.

#### Establishment of ccRCC and Normal Kidney Organoids

Fresh or thawed PDTFs were enzymatically digested in 2 mg/mL Collagenase P (Merck) in Dulbecco’s Modified Eagle Medium (DMEM) (Thermo Fisher Scientific) for one hour at 37°C in a thermal shaker. Digestion was monitored every 10 minutes after the first 30 minutes to prevent over-digestion. The digested tissue suspensions were sequentially filtered through 100-μm and 40-μm cell strainers (pluriSelect), centrifuged to pellet the cells, and resuspended in red blood cell lysis buffer (Invitrogen) for 3 minutes to remove red blood cells. The final cell pellet was washed with phosphate-buffered saline (PBS), and the viable cell yield was determined using Trypan Blue (Invitrogen) staining and manual counting. Single-cell suspensions were processed according to the starting material and target cell population, as described below:

#### Bulk normal or tumour organoids

Cell suspensions were pelleted and resuspended in 75% Matrigel (Corning) at 1 million cells per 100 μL. The Matrigel-cell mixture was plated as droplets, with plates inverted and incubated at 37°C for 20 minutes to solidify before the addition of complete organoid medium.

#### CD24+ CD13+ VCAM1+ normal organoids

For the establishment of normal proximal tubule-derived VCAM1+ organoids, single-cell suspensions were pelleted and resuspended in staining solution at a concentration of 1 million cells per 50 μL. The following antibodies were used for fluorescence-activated cell sorting (FACS): anti-CD24-BV421 (BioLegend, 1:20), anti-CD13-PE (BioLegend, 1:20), and anti-VCAM1-APC (Miltenyi, 1:10). After 30 minutes of incubation in the dark, cells were washed and resuspended in FACS buffer (2 mM EDTA, 0.5% bovine serum albumin (BSA) in PBS) with live-dead dye. Autofluorescence was excluded using an irrelevant fluorescence channel, and fluorescence-minus-one controls were used for proper gating. CD24+ CD13+ cells were first gated, followed by VCAM1⁺ selection, and sorted using a FACSAria Fusion sorter (BD Biosciences). The sorted cell population was pelleted and resuspended in a 75% Matrigel-medium mixture at a concentration of 1 million cells per 100 μL, plated as droplets, and incubated at 37°C for 20 minutes before the addition of complete medium.

#### CA9+ tumour organoids

For the establishment of tumour-derived CA9+ organoids, single-cell suspensions were pelleted and resuspended in staining solution at a concentration of 1 million cells per 50 μL. The following antibodies were used: anti-CA9-PE (Miltenyi, 1:50), anti-CD45-APC (BioLegend, 1:50), and DAPI (Thermo Fisher Scientific). After 30 minutes of incubation in the dark, cells were washed and resuspended in FACS buffer. Live cells were gated, and CA9+ (tumour cells) and CD45+ (tumour-infiltrating lymphocytes, TILs) populations were sorted separately using a FACSAria Fusion sorter (BD Biosciences). Sorted CA9+ tumour cells were seeded in a 75% Matrigel-medium mixture at a concentration of 1 million cells per 100 μL, plated as droplets, and incubated at 37°C for 20 minutes before the addition of complete medium. CD45+ TILs were cryopreserved. As an alternative to FACS-based positive selection, Magnetic-Activated Cell Sorting (MACS)-based negative selection was performed using the CD45 MicroBead Kit (Miltenyi), following the manufacturer’s protocol. Both bead-bound (CD45+) and flow-through (CD45- tumour cell-enriched) fractions were collected. CD45+ TILs were cryopreserved, while CD45- tumour-enriched cells were seeded in Matrigel as described above.

#### Maintenance of ccRCC and Normal Kidney Organoids

Cells isolated from normal and tumour tissues were embedded in Matrigel droplets and cultured in normal organoid growth medium or tumour organoid growth medium (**Table S8**), as previously described^65,66^. Tumour organoids were maintained in a hypoxia (1% O_2_) incubator. Organoids were passaged upon reaching confluence or when no further growth was observed. Normal PDOs typically required approximately a week to recover after the initial passage, while tumour PDOs exhibited variable growth rates. For organoid dissociation during passaging, TrypLE Express (Gibco) was used, with digestion times differing between normal and tumour organoids. Normal organoids required longer digestion times (7–10 minutes), whereas tumour organoids dissociated more rapidly, and they were sensitive to prolonged digestion, requiring careful monitoring to prevent over-digestion. To preserve organoid integrity, particularly in slow-growing or fragile lines, Dispase II (Thermo Fisher Scientific) was used to digest Matrigel while maintaining organoid structure selectively. The splitting ratio was adjusted based on the proliferation rate of each line: rapidly proliferating organoids were split at a 1:3 ratio, whereas slow-growing organoids were split at a 1:1 ratio or higher (to concentrate rather than dilute the culture). For cryopreservation, organoids were harvested following the same procedures as for passaging. Cells were resuspended in freezing medium (90% FBS + 10% DMSO) and cryopreserved using a programmed cooling system to −80°C before being transferred to liquid nitrogen for long-term storage. For thawing, frozen cryovials were warmed in a 37°C water bath until a small ice pellet remained. The thawed suspension was transferred to a 15-mL Falcon tube, and 10 mL of warm Advanced DMEM medium (Thermo Fisher Scientific) was added dropwise while gently flipping the tube to prevent osmotic shock. Cells were pelleted by centrifugation and resuspended at a high density (matching the initial cryopreservation density) to optimise recovery and ensure efficient regrowth.

#### Characterisation of Tumour Organoids

Tumour organoid outgrowth was validated through genomic mutation analysis and/or CA9 staining to confirm tumour cell identity. When the primary tumour’s genomic profile was available, a pair of custom-designed primers was used to amplify the mutation-containing region, followed by Sanger sequencing or amplicon sequencing. The relative tumour purity of the organoid cultures was then estimated using SnapGene or Integrative Genomics Viewer (IGV). In cases where no primary tumour sequencing data were available, *VHL* amplicon sequencing was performed as an alternative. Additionally, organoids were harvested and subjected to CA9 staining, as described in the previous section, to quantify the proportion of CA9-positive tumour cells using a ZE5 flow cytometry analyser (Bio-Rad). A verified normal organoid culture was included as a negative control to ensure specificity in CA9-based validation.

#### DNA & RNA Extraction

For DNA and RNA extraction used in omics profiling, the AllPrep DNA/RNA Mini Kit (Qiagen) was used. Briefly, a 1–2 mm³ piece of frozen tissue was excised and added to 600 μL of lysis buffer, followed by homogenisation using a TissueLyser (Qiagen) and further processing through a QIAshredder (Qiagen) spin column to collect the cell lysate. For cultured organoids and cells, cell pellets were lysed in 600 μL of lysis buffer and similarly processed using a QIAshredder (Qiagen) spin column to obtain the lysate. Nucleic acid purification was performed using binding columns according to the manufacturer’s protocol. For DNA extraction intended for genotyping and Sanger sequencing, the Monarch Spin gDNA Extraction Kit (NEB) was used following the manufacturer’s guidelines. DNA and RNA concentrations were measured using a Qubit fluorometer (Thermo Fisher Scientific) or a Nanodrop spectrophotometer (Thermo Fisher Scientific). RNA integrity and quality were assessed using either a TapeStation (Agilent) or a Bioanalyzer (Agilent).

#### Organoid formation assay

Organoids were collected and enzymatically dissociated using dispase II (Thermo Fisher Scientific) for 30 minutes, followed by Accutase (Thermo Fisher Scientific) for 15 minutes, to obtain a single-cell suspension. Single cells were then resuspended in Matrigel at a concentration of 1,000 cells per μL. Organoid formation efficiency was monitored via ECHO Revolution microscope with brightfield channel over a period of 20 days.

#### Haematoxylin & Eosin Staining

Matrigel-embedded organoids were dissociated using Gentle Cell Dissociation Reagent (Stem Cell Technologies) at 4°C on a shaker until the Matrigel was fully dissociated. The harvested organoids were fixed in 10% neutral buffered formalin (NBF) for 1.5 hours at 4°C on a roller. Following fixation, organoids were washed with PBS, embedded in agarose in a 1.5 mL tube, and processed into paraffin blocks for sectioning. Haematoxylin and eosin (H&E) staining were performed at the Experimental Histopathology (EHP) facility at the Francis Crick Institute with standard procedures.

#### Micronuclei staining

Organoids were enzymatically dissociated into single cells using dispase II (Thermo Fisher Scientific) for 30 min, followed by Accutase (Thermo Fisher Scientific) for 15 min. The resulting single-cell suspension was fixed in 4% paraformaldehyde (PFA) for 20 min at room temperature, washed with PBS, and stained with 0.1 µg/mL DAPI for 10 min. After additional PBS washes, cells were mounted on glass slides and imaged using ECHO Revolution microscope with DAPI channel.

#### Genetic Manipulation in Organoids

Electroporation using the Lonza system was employed for fast and efficient gene editing. Organoids were collected and enzymatically dissociated using dispase II (Thermo Fisher Scientific) for 30 minutes, followed by Accutase (Thermo Fisher Scientific) for 15 minutes, to obtain a single-cell suspension. The CRISPR ribonucleoprotein (RNP) complex was assembled by incubating 2 μL of 100 μM gRNA solution (Integrated DNA Technologies (IDT)) with 2 μL of 50 μM Alt-R HiFi-Cas9 protein (IDT) for 30–45 minutes. A total of one million single cells were pelleted and resuspended in 100 μL electroporation buffer (Amaxa P3 Primary Cell Solution, Lonza), prepared by adding 18 μL of Supplement to 82 μL P3 Solution. The CRISPR RNP complex was then added to the cell suspension, which was transferred into a Lonza 100 μL cuvette and electroporated using the Amaxa 4D Nucleofector (Lonza) with the ‘Primary Cell P3’ program and ‘CM-138’ pulse code. Immediately after electroporation, 1000 μL of pre-warmed culture medium was added to the reaction mix and incubated for 5–10 minutes before seeding the cells into 200 μL Matrigel in four wells of a 12-well plate. To assess knockout efficiency, one well of organoid culture was harvested, followed by DNA extraction and *VHL* Sanger sequencing or amplicon sequencing. Sanger sequencing results were analysed with ICE (Inference of CRISPR Edits) to determine efficiency^67^. To obtain a pure population of engineered organoids, manual picking and genotyping were used for enrichment. Organoid cultures were dissociated with Dispase II for 30 minutes to isolate single organoids, which were then resuspended in PBS with 1% BSA. Individual organoids were manually picked under a brightfield microscope using a 10 μL pipette and transferred into 50 μL TrypLE Express for 3 minutes of dissociation. The resulting cell suspensions were pelleted and seeded into a 48-well plate for expansion. Expanded organoids were harvested for *VHL* amplicon sequencing to determine mutation frequency. Organoids exhibiting a mutation frequency higher than 90% were retained and used as the pure engineered population for subsequent experiments.

#### Amplicon Sequencing

Genomic DNA was extracted, and 50 ng was used for multiplex PCR with three pairs of primers designed to specifically amplify the three exons of *VHL* while incorporating a universal linker. A second PCR was performed using Nextera library sequencing primers (S5/7-ME), which contained sequences complementary to the linker. Following the second PCR, the products were further amplified using Nextera library adaptor-index primers, enabling sample multiplexing. The final amplicon libraries were sequenced using the MiSeq v2 250 bp paired end (PE) platform. Sequencing output data were mapped against the reference genome GRCh38 with Bowtie2 (v 2.3.4.2), and *VHL* mutations were manually inspected using IGV. Variant allele frequencies (VAFs) were calculated accordingly to assess the presence and proportion of *VHL* mutations.

#### *VHL* Methylation Sequencing

*VHL* promoter methylation was assessed following bisulphite conversion and purification of 200 ng of genomic DNA using the EZ DNA Methylation-Direct kit (Zymo Research). The bisulphite-treated DNA was then subjected to methylation-specific PCR to amplify the *VHL* promoter region. To confirm methylation status, Sanger sequencing was performed on the MSP amplicons. Methylation was determined by comparing tumour-derived samples with normal renal tissue, specifically assessing methylation-protected CpG sites, where cytosines remained unconverted due to methylation.

#### ccRCC Driver Panel Sequencing and Analysis

Library preparation and sequencing

Targeted sequencing of ccRCC driver genes was performed using the custom version 7 TRACERx panel (Agilent), as previously described in the original TRACERx studies^9,19^ . Library preparation involved target enrichment using a capture bait set (Agilent) covering 130 cancer driver genes along with a SNP backbone for copy number and purity estimation. The captured libraries were then sequenced using Illumina HiSeq 2500 and NextSeq 500 platforms at the Genomics Facility of the Francis Crick Institute.

#### Data processing

Raw FASTQ files from germline samples (blood and/or normal organoids) and multiregional tumour samples (primary tumour and/or tumour organoids) were processed using a customised in-house bioinformatics pipeline benchmarked for panel sequencing analysis. The Nextflow Sarek pipeline was modified to support multiregional panel sequencing data, using a custom BED file (Panel_v7_hg38.bed). Raw sequencing reads were aligned to the GRCh38 human genome reference using BWA (v.0.7.17-r1188)^68^. Variant calling was performed using FreeBayes (v.1.3.2)^69^, and variant annotation was conducted with VEP (v.106.1)^70^. CNV calling, tumour purity estimation, and ploidy assessment were carried out using ASCAT (v.25.2)^71^ and FACETS (v.0.6.2)^72^. Tumour mutation and CNV profiles were visualised as an oncoplot. Tumour phylogenetic trees (clone trees) were inferred manually based on clonality assessments and mutational frequency distributions.

#### Low Pass Whole Genome Sequencing

Genomic DNA was extracted, and library preparation was conducted using the Nextera Flex protocol, followed by sequencing at 0.5-1x coverage. Copy number calling and karyotyping were performed as previously described using QDNAseq (v1.38.0)^73^. Briefly, the genome was segmented into 100 kb bins, and read counts were calculated and corrected for GC content and mapability biases. Log2-transformed copy number ratios were derived by comparing observed read counts to expected counts, and copy number alterations were visualised along chromosomal coordinates.

#### Bulk RNA Sequencing and Analysis

Library preparation and sequencing

RNA-seq library preparation and sequencing were performed on organoid samples with sufficient RNA quality (RNA integrity (RIN) score ≥ 5, assessed using TapeStation 4200 or Bioanalyzer). High-quality RNA samples were processed by the Genomics Facility at the Francis Crick Institute. RNA input was normalised to 100 ng, and libraries were prepared using the Watchmaker RNA Library Prep Kit with Polaris™ Depletion (Watchmaker Genomics), following the manufacturer’s protocol. Ribosomal RNA (rRNA) and globin transcripts were depleted before library construction to enrich for mRNA. Libraries were indexed with unique dual indexes (TruSeq UD Indexes, 20022371) and PCR-amplified for 15 cycles. Library quality and fragment size distributions were assessed using TapeStation 4200 (Agilent). Final libraries were pooled and subjected to paired-end sequencing on an Illumina HiSeq4000 or NovaSeq 6000 platform.

Data processing

Raw FASTQ files were processed using the Nextflow RNA-seq pipeline (v.3.6). Briefly, sequencing quality was assessed using FastQC (v.0.11.9). Alignment to the GRCh38 human genome and gene expression quantification were performed using STAR-RSEM (v.2.7.6a, v.1.3.1)^74^. The resulting raw gene count table was used for downstream analysis. A DESeq2^75^ object was constructed, and variance-stabilizing transformation was applied for normalisation. Unless otherwise specified, these normalised counts were used in subsequent analyses.

### Bulk ATAC Sequencing and Analysis

Library preparation and sequencing

Organoids were dissociated into single cells as described above. 60,000 single cells were cryopreserved in 50 μL BAM Banker Cryopreservative (Fisher Scientific, cat. no. NC9582225). ATAC-seq library preparation was performed following a published ATAC-seq protocol^76^. Briefly, 60,000 cryopreserved single cells were thawed and used as input per reaction. Nuclei were extracted by incubating cells in ATAC-seq Lysis Buffer for exactly 3 minutes on ice. The isolated nuclei were then incubated with the Transposition Mix (Illumina) at 37°C for 30 minutes in a thermomixer at 1,000 rpm, generating transposed DNA fragments. These fragments were purified and eluted in nuclease-free water using the DNA Clean & Concentrator-5 kit (Zymo Research). Purified transposed fragments were barcoded and PCR-amplified using a Nextera library structure, with the exact number of amplification cycles determined using the NEBNext Library Quant Kit (NEB) for Illumina, following the manufacturer’s instructions. Library quality and fragment size distributions were assessed using TapeStation 4200 (Agilent). Final libraries were sequenced at a target depth of 25 million PE-100 bp reads per sample on the Illumina NovaSeq 6000 at the Genomics Facility at the Francis Crick Institute.

Data processing

Raw FASTQ files were processed using the Nextflow ATAC-seq pipeline (v.1.2.1). Sequencing quality was assessed using FastQC (v.0.11.9). Reads were aligned to the GRCh38 human genome using BWA (v.0.7.17-r1188)^68^, and coverage normalisation was performed with deepTools (v.3.4.3)^77^. BigWig files generated from the normalised coverage data were directly loaded into IGV for visualisation. JUNB peak accessibility heatmaps were generated with deepTools computeMatrix and plotHeatmap functions using a JUNB ChIP-seq peak bed derived from A549 cells (ENCODE ID: ENCFF971EYD)^78^.

### Single Cell RNA Sequencing and Analysis

Library preparation and sequencing

Cultured organoids were dissociated into single-cell suspensions as described above, and up to 1 million single cells per sample were fixed using the Evercode Fixation v3 kit (Parse Bioscience) and cryopreserved at −80°C for long-term storage, following the manufacturer’s protocol. A total of 12 fixed and cryopreserved samples were collected and processed in a single multiplexed barcoded single cell sequencing run. For cDNA library preparation, the Evercode WT Mini v3 kit (Parse Bioscience) was used, targeting 1,650 cells per sample in accordance with the manufacturer’s instructions. The quality and fragment size distribution of the final library were assessed using the TapeStation 4200, and libraries were sequenced at an average depth of 60,000 PE100 reads per cell on the Illumina NovaSeq 6000 at the Genomics Facility at the Francis Crick Institute.

Data processing

Raw FASTQ files were processed using the Parse Bioscience pipeline, split-pipe (v1.2.1), with default parameters to align sequencing reads to the GRCh38 human genome, demultiplex samples, and quantify gene expression as a count matrix. The resulting count matrices for each sample were loaded into Seurat. To ensure high-quality data, low-quality cells and possible doublets (fewer than 200 RNA features, more than 10,000 RNA features, or mitochondrial RNA content exceeding 30%) were removed. Seurat’s standard pipeline was then applied for clustering and visualisation, including NormalizeData, ScaleData, RunPCA, FindNeighbors (using the top 30 PCA dimensions), FindClusters (resolution = 0.5), and RunUMAP functions. Gene expression was visualised using DotPlot with scaling enabled. For copy number variation analysis, InferCNV (v.1.3.3)^79^ was applied, using the normal organoid line K1030N1_GN1 as a reference.

### Cohort Curation of ccRCC Bulk RNA-seq Datasets

**TCGA-KIRC cohort:** Raw RNA-Seq counts from 538 TCGA-KIRC primary tumour samples were downloaded using the TCGAbiolinks R package (v.2.20.0)^80^, with the following parameters: data.category = “Transcriptome Profiling”, data.type = “Gene Expression Quantification”, experimental.strategy = “RNA-Seq”, workflow.type = “STAR-Counts”. The count table was filtered to include only protein-coding genes. Metadata, including sample annotations and genotypes, were obtained from cBioPortal. Overlapping expression data with metadata resulted in 507 tumour samples. The gene expression count table and metadata were used to construct a DESeq2 object, which was normalised using variance stabilising transformation (VST)^75^. To subset samples with *VHL* inactivation, cases were selected based on at least one of the following criteria: 1) the presence of a somatic mutation in *VHL*, 2) promoter methylation at three *VHL* promoter CpG sites (cg16869108 ≥ 0.8, cg20916523 ≥ 0.6, cg25539131 ≥ 0.8), or 3) mRNA expression z-scores relative to normal samples ≤ −1. Cases with *PBRM1* or *BAP1* mutations were further selected, retaining a total of 136 samples for analysis.

**TRACERx Renal cohort:** Raw RNA-Seq counts, metadata, and genotypes from 224 TRACERx Renal samples were sourced in-house^27^, processed into a DESeq2 object, and normalised using VST. Applying the *VHL*, *PBRM1*, and *BAP1* alteration status filtering criteria retained 101 samples.

**Tumourgraft (TG) cohort:** TPM (Transcripts Per Million) data and metadata for 71 tumour xenograft samples were obtained from the supplementary data of Elias et al.^28^. After filtering for *VHL* mutations and *PBRM1* and/or *BAP1* mutations, 42 samples were retained.

### RNA-seq Signature Score Estimation

To estimate signature scores across the three bulk RNA-seq datasets, single-sample Gene Set Enrichment Analysis (ssGSEA) was performed using the R package GSVA (v.1.48.3)^81^. Normalised expression values, either TPMs or VST-transformed counts, were used as input, with default parameters applied as recommended. The pathway analysis in this study included 23 Hallmark gene sets from the Molecular Signatures Database (MSigDB)^82^, 13 metaprograms (MPs) derived from pan-cancer scRNA-seq data^83^, two proximal tubule marker sets from the Human Protein Atlas^84^, two published gene signatures (HIF_meta_gene^85^ and Epithelial_Sig_Kidney^86^), and one in-house curated PT_VCAM1 signature (**Table S4**). For signature score calculation in scRNA-seq data, the AddModuleScore_UCell function in the UCell R package^87^ was used.

### TME cell type estimation

Bulk RNA-seq data from TCGA-KIRC and TRACERx RNA cohorts were used as input for deconvolution by the consensusTME R package^38^. The parameter “statMethod” was set to “ssGSEA” and “cancer” was set to “KIRC”.

### Curation of ccRCC scRNA-seq Datasets

Single-cell RNA-seq data from six studies^31–36^ were downloaded either as CellRanger output files or Seurat objects, depending on the formats provided in the original publications. Metadata at both the sample and single-cell levels were collected and processed to generate a unified dataset, incorporating key annotations including Disease_type, Tissue_type, Patient_id, Sample_origin, Stage, Treatment, Platform, scORsn, Lib_type, Enrichment, Lineage, Initial_cluster, Initial_annotation, InferCNV_status, and Genotype. Using the Seurat (v5.0.1)^88^ R package, unified Seurat objects were constructed by integrating the expression matrix and metadata. To remove low-quality cells and doublets, cells in which mitochondrial reads exceeded 30% of total reads were filtered out, as kidney epithelial and ccRCC cells are known to exhibit high mitochondrial RNA content. Cells with fewer than 200 RNA features or more than 10,000 RNA features were also excluded. Gene features were filtered to only include protein-coding genes. A total of 589,212 high-quality cells from different studies were then integrated using canonical correlation analysis (CCA), reciprocal PCA (RPCA), and Harmony^89^ within Seurat’s built-in function IntegrateLayers, applied to PCA dimensions. CCA integration was selected for further annotation. Dimensionality reduction and clustering were performed using UMAP on the integrated dimensions. Cell type annotation was conducted using a panel of known marker genes. Non-immune clusters, including Fibroblasts, Endothelial cells, and Tumour_kidney_cells (Epithelial cells), were subsequently subset and re-clustered following the same integration and annotation pipeline. To specifically identify ccRCC tumour cells, InferCNV^79^ was applied to the non-immune clusters, using fibroblasts or endothelial cells as normal references. Mean residual expression values for genes located in the chr3p region (chr3:8,100,001-54,400,000, spanning from chr3p25 to chr3p21) were computed for each cell. Cells with a mean residual expression below −0.05, indicative of 3p loss, were classified as tumour cells. After applying this classification across all six studies, a total of 119,036 tumour cells were retained for downstream analyses.

| Cell Type | Markers |
| --- | --- |
| B cells | "CD19", "MS4A1", "CD79A" |
| Endothelial cells | "ADGRL4", "EGFL7", "PLVAP" |
| Fibroblast | "COL6A2", "LUM", "MFAP5" |
| Myeloid cells | "CD14", "FCGR3A", "ITGAM" |
| Plasma cells | "IGKC", "IGHG1", "SDC1", "TNFRSF17" |
| T cells | "PTPRC", "CD3E" |
| Tumour_kidney_cells | "CA9", "CP", "CUBN", "PDZK1IP1" |

### scRNA-seq and Bulk RNA-seq Differential Analysis

For differential expression analysis in scRNA-seq data from both clinical specimens and PDOs, the FindMarkers function in Seurat was applied, with ident.1 set to the population of interest and ident.2 set as the control group. The Wilcoxon Rank Sum test was used as the statistical method (test.use = “wilcox”) unless otherwise specified. Differentially expressed genes were filtered using an adjusted p-value threshold of ≤0.05. For bulk RNA-seq differential expression analysis, DESeq2 was applied to the raw gene count data. The resulting differentially expressed gene lists were used for visualisation in Volcano plots (fold change vs. −log p-value) and MA plots (log difference vs. log average) generated using ggplot2 (v.3.5.1). For MA plots, lfcshrink values were used, as recommended in the DESeq2 manual, to improve visualisation by shrinking log-fold changes for low-expression genes. Gene set enrichment analysis (GSEA) was performed using the fgsea R package (v.1.28.0)^90^, with the MSigDB hallmark dataset used as the reference pathway collection. The resulting normalised enrichment scores (NES) and adjusted p-values were visualised using ggplot2 (v.3.5.1).

### Cell-cell interaction inference

To infer the probability of cell-cell interactions in scRNA-seq datasets, we used the R package CellChat (v2.1.0)^91^ with the ligand-receptor database CellChatDB (v2)^92^. Briefly, we separated *PBRM1*-driven tumours and *BAP1*-driven tumours as two Seurat objects and then converted them into CellChat objects. We then applied the ‘identifyOverExpressedGenes’ and ‘identifyOverExpressedInteractions’ with the default settings, which resulted in around 2000 highly variable ligand-receptor pairs in both groups for further signalling inference. We calculated the communication probability/strength with ‘computeCommunProb’ with ‘triMean’ method and filtered out communications for cell groups with cells less than 10. Then we applied ‘computeCommunProbPathway’ and ‘aggregateNet’ to aggregate the pathway-level interaction network. Finally, we identify signalling roles of cell types with the ‘netAnalysis_computeCentrality’ function and visualise the results with the function ‘netAnalysis_signalingRole_scatter’.

### scRNA-seq Regulon Analysis

To infer gene regulatory networks, the SCENIC pipeline was implemented using the pySCENIC command-line interface (v.0.12.1) within a conda environment^45^ . The analysis was performed on the RNA assay of the PDO scRNA-seq dataset. For the initial step, the GRNBoost2 algorithm was used, as recommended in the SCENIC manual for large-scale datasets. A curated list of transcription factors (TFs) was downloaded from the pySCENIC website and used as input. Regulon prediction and cellular enrichment were performed with default parameters using the RcisTarget databases provided on the pySCENIC website (hg38_10kbp_up_10kbp_down_full_tx_v10_clust.genes_vs_motifs.rankings.feather). The resulting AUCell matrix of regulon activity was then used for UMAP clustering. Due to the stochastic nature of the SCENIC pipeline, as noted in the original publication, the full analysis was repeated three times to improve reproducibility. Regulons that consistently appeared across all three runs were retained. To prioritise key regulons for Figure 3A and Figure S3A, the distribution of regulon scores across samples in all three runs was examined, and those with stable and consistent distributions were selected for heatmap presentation.

### ATAC-seq Differential Analysis and Transcriptional Factor Footprinting

ATAC-seq peak calling was performed using MACS3 (v3.0.3)^93^ with the parameters --nomodel --shift 100 --extsize 200. Peak score matrices were generated with the DiffBind R package (v3.12.0)^94^, followed by peak annotation using the annotatePeak function in ChIPseeker (v1.38.0)^95^. Transcription factor (TF) footprinting analysis was conducted using TOBIAS (v0.17.2)^96^. Briefly, sequence bias introduced by Tn5 transposase cleavage was corrected with the ATACorrect function, and footprint scores were computed using the FootprintScores function. The BINDetect function was then applied to infer TF binding activity within individual samples and to assess differential binding across samples. Motif reference files were obtained from the JASPAR 2024 CORE database (https://jaspar.elixir.no/download/data/2024/CORE).

## QUANTIFICATION AND STATISTICAL ANALYSIS

For all the statistical tests, the significance level was fixed at 0.05 unless otherwise specified. Sample sizes (n) and tests used are specified either by showing in the figures and/or by an indication in the figure legends. Statistical tests were performed in either GraphPad Prism or RStudio.

